# Genuine cross-frequency coupling networks in human resting-state electrophysiological recordings

**DOI:** 10.1101/547638

**Authors:** Felix Siebenhühner, Sheng H Wang, Gabriele Arnulfo, Anna Lampinen, Lino Nobili, J. Matias Palva, Satu Palva

## Abstract

Phase synchronization of neuronal oscillations in specific frequency bands coordinates anatomically distributed neuronal processing and communication. Typically, oscillations and synchronization take place concurrently in many distinct frequencies, which serve separate computational roles in cognitive functions. While within-frequency phase synchronization has been studied extensively, less is known about the mechanisms that govern neuronal processing distributed across frequencies and brain regions. Such integration of processing between frequencies could be achieved via cross-frequency coupling (CFC), either by phase-amplitude coupling (PAC) or by *n:m*-cross-frequency phase synchrony (CFS). So far, studies have mostly focused on local CFC in individual brain regions, whereas the presence and functional organization of CFC between brain areas have remained largely unknown. We posit that inter-areal CFC may be essential for large-scale coordination of neuronal activity and investigate here whether genuine CFC networks are present in human resting-state brain activity.

To assess the functional organization of CFC networks, we identified brain-wide CFC networks at meso-scale resolution from stereo-electroencephalography (SEEG) and at macro-scale resolution from source-reconstructed magnetoencephalography (MEG) data. We developed a novel graph-theoretical method to distinguish genuine CFC from spurious CFC that may arise from non-sinusoidal signals ubiquitous in neuronal activity. We show that genuine inter-areal CFC is present in human resting-state activity in both MEG and SEEG data. Both CFS and PAC networks coupled theta and alpha oscillations with higher frequencies in large-scale networks connecting anterior and posterior brain regions. CFS and PAC networks had distinct spectral patterns and opposing distribution of low- and high frequency network hubs, implying that they constitute distinct CFC mechanisms. The strength of CFS networks was also predictive of cognitive performance in a separate neuropsychological assessment. In conclusion, these results provide evidence for inter-areal CFS and PAC being two distinct mechanisms for coupling oscillations across frequencies in large-scale brain networks.

## Introduction

Human electrophysiological activity is characterized by neuronal oscillations, *i.e.*, rhythmic excitability fluctuations in a wide range of frequencies, at least from 0.01 to over 150 Hz. Synchronization of these oscillations across brain areas coordinates and regulates anatomically distributed neuronal processing [1, 2]. In humans, large-scale oscillatory networks in several frequency bands characterize magnetoencephalography (MEG), electroencephalography (EEG), and stereo-EEG (SEEG) data during resting state (RS) activity [3-8] and in many cognitive functions [9-13]. Inter-areal synchronization of alpha (α, 7-14 Hz) and beta (β, 14-30 Hz) oscillations in humans and non-human primates, respectively, is thought to regulate top-down or feed-back communication [14-19]. In contrast, both β and gamma-band (γ, 30-100 Hz) oscillations and synchronization have been associated with bottom-up sensory processing and representation of object-specific sensory information [15, 20-22], and β oscillations are also with sensorimotor processing [23, 24]. Overall, brain-wide oscillation networks in multiple frequencies are proposed to be the core of cognition [10, 11, 25-27]. Also, human brain activity at rest is characterized resting-state networks (RSNs) first identified with fMRI [28, 29]. Oscillatory RSNs observed in electrophysiological measurements are organized in a partially similar fashion as the RSNs observed with fMRI [3, 30] as well as into a modular structure at the whole-brain connectome level [31]. It has been proposed that RSNs form the basis of task-state large-scale networks [8, 32].

The interplay between oscillations at distinct frequencies is thought to be regulated via two forms of cross-frequency coupling (CFC): phase-amplitude coupling (PAC) [9, 33-37] and cross-frequency phase synchrony (CFS) [9, 38-41], also known as *n*:*m*-phase synchrony [38]. PAC indicates the correlation between the amplitude envelope of a faster oscillation and the phase of a slower oscillation, whereas CFS is a form of phase synchronization defined by a non-random phase difference between oscillations having an integer *n:m* frequency ratio (Fig 1a). During task performance, PAC has been suggested to reflect the regulation of sensory information processing in β and γ-frequencies by excitability fluctuations imposed by θ and α oscillations [9, 10, 34, 36, 37, 42]. A large number of studies have identified local PAC, *i.e.*, PAC observed between different frequency bands of the same signal, between the phases of slower oscillations in delta (δ, 1–3 Hz), theta (θ, 3–7 Hz) and α frequency bands and the amplitude of γ oscillations in local field potentials (LFPs) in rats [43-47] and in human intracranial EEG [48-51] and MEG data [52-56]. Such local PAC has cortex-wide spatial modes akin to RSNs [53]. Unlike PAC, CFS enables temporally precise coordination of neuronal processing by establishing systematic spike-timing relationships among possibly functionally distinct oscillatory assemblies and hence has been suggested to serve functional integration and coordination across within-frequency synchronized large-scale networks [39, 41, 57]. Local CFS has been observed in human MEG and EEG data during rest [39, 58] and during attentional and working memory (WM) tasks [39, 40, 59] as well as in LFPs in the rat hippocampus [45].

**Fig 1.**
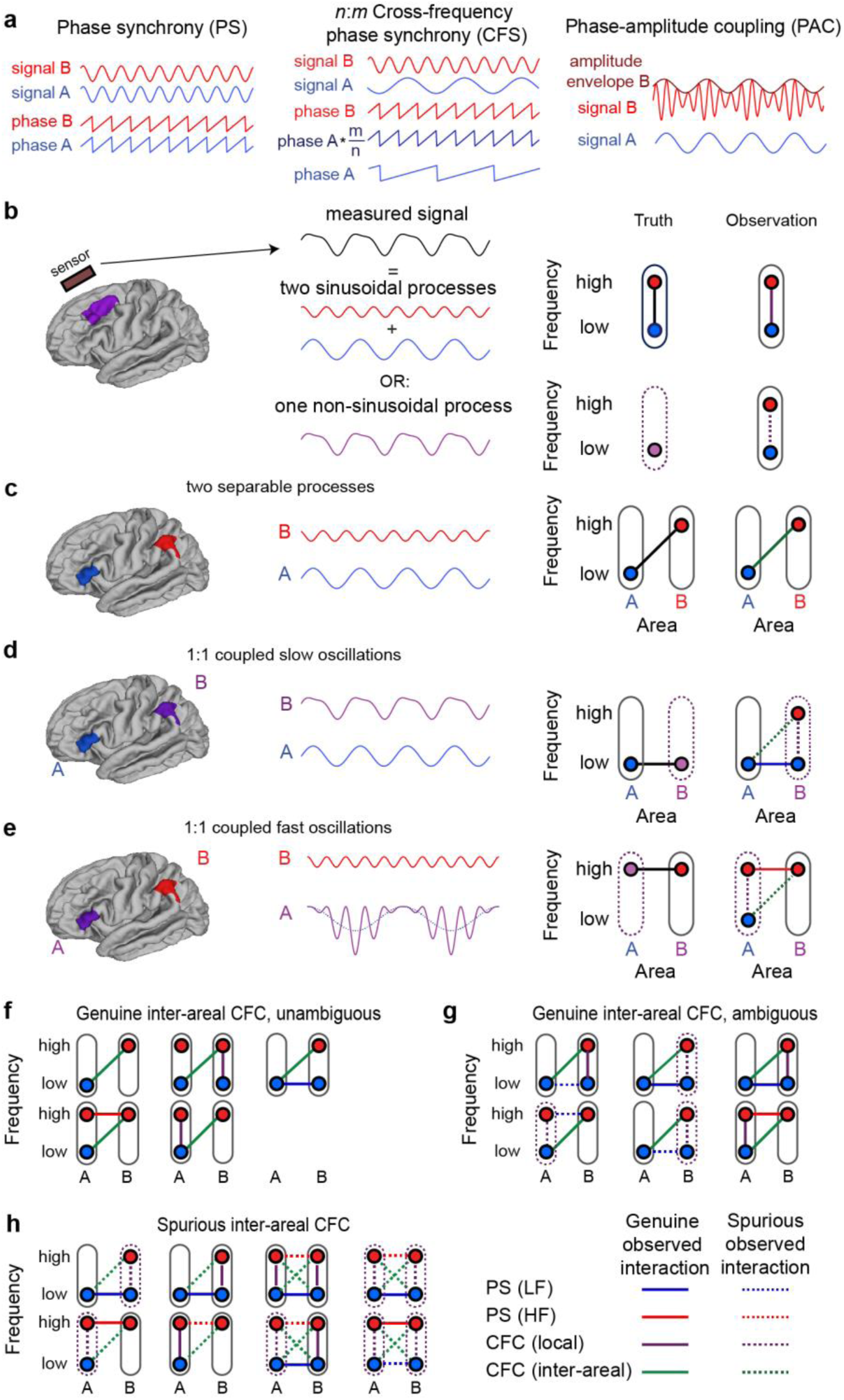
Schematics of for identifying genuine and spurious inter-areal CFC. **a)** Schematic illustration of phase synchrony (PS), LF:HF (*n*:*m*) cross-frequency phase synchrony (CFS) and LF:HF phase-amplitude coupling (PAC). In PS, two spatially distant processes oscillating at the same frequency exhibit a (statistically) constant phase relationship. In CFS, a constant *n*:*m*-phase relationship exists between two processes at frequencies LF and HF, so that LF:HF = *n*:*m*. In PAC, the amplitude of the high-frequency signal is correlated with the phase of the low-frequency signal. These processes can either take place in the same region (local CFC) or between two regions (inter-areal CFC). **b)** Observations of local CFC can be either genuine or spurious. A measured signal from a single sensor or electrode can either be the sum of two (statistically) sinusoidal processes oscillating at distinct frequencies or a single non-sinusoidal process and these possibilities are difficult to dissociate from the single signal. Local CFC can be observed in both cases because of filter artefacts produced by non-sinusoidal signals. **c)** Genuine inter-areal CFC between two spatially distant sinusoidal processes A and B. **d)** An example of spurious observation of inter-areal CFC. Process A is sinusoidal, but B is non-sinusoidal and causes spurious local CFC to be observed at location B, as shown in b). If A and B are connected by low-frequency PS, also spurious *inter-areal* CFC will be observed between A and B. This spurious observed inter-areal CFC forms a “triangle motif” with PS and the spurious CFC couplings. **e)** Example of spurious observation of inter-areal CFC where process B is sinusoidal, but A has a non-zero-mean and spurious local CFC will be observed at location A. Again, if A and B are connected by HF PS, also spurious inter-areal CFC will be observed between A and B, again forming a triangle motif. **f)** Constellations of observations that unambiguously indicate genuine CFC between regions A and B. In none of the cases there is a triangle motif of PS and local CFC. **g**) Constellations of observations with ambiguous finding of inter-areal CFC between regions A and B. Although here inter-areal CFC is genuine, it is part of a triangle motif formed with PS and (true or spurious) local CFC. Such constellations cannot be distinguished by our graph-theory based method from spurious inter-areal CFC. **h**) Constellations with spurious inter-areal CFC. These include the two constellations from d) and e) in the left column and other possible constellations, including those where there is spurious local CFC at both locations (right column). In all cases, the spurious inter-areal CFC is part of a triangle motif.

However, observations of CFS have remained scarcer than those of PAC and it has remained unclear whether CFS and PAC are even distinct phenomena or simply different reflections of a single CFC phenomenon. Some studies have also identified inter-areal PAC or CFS between few pre-selected regions of interest between MEG/EEG sensors [39, 40] or between brain regions in source-reconstructed MEG/EEG data [41, 53, 56] or intracranial data [60]. However, only few studies [39, 41] have identified CFC in cortex-wide networks that form the basis of cognitive functions. Therefore, it has remained largely unknown if CFC can couple oscillations across frequencies in large-scale brain networks and, if so, what is the functional organization of these networks, their similarities to PS networks, and their relevance for cognitive performance and abilities. Furthermore, recent research has suggested that estimates of CFC may be inflated by false positive couplings arising from non-sinusoidal and non-zero mean signals. False positives are caused by the artificial higher-frequency components produced when non-sinusoidal signals are filtered into narrow bands [61-68], as well as by artificial lower-frequency components arising from filtering of non-zero-mean waveforms [69]. Because non-sinusoidal and non-zero-mean waveforms are ubiquitous in electrophysiological signals, their filter artefacts constitute a significant confounder to CFS and PAC estimation and question the validity of prior CFC observations. As CFC is thought to be central in the coordination of processing across frequencies, it is crucial to establish if genuine neuronal CFC can be observed in neuronal activity in the first place.

The fundamental assumption in CFC is that it indicates an interaction between *two* distinct neuronal processes in two frequency bands. Conversely, the notion of artificial CFC arising from non-sinusoidal signal properties relies on the assumption that a neuronal process exclusively in a *single* frequency band generates the observed signal. Approaches based on waveform analysis [63, 68, 70] and appropriate control analyses [69] have been proposed to reduce the artefactual connections. Nevertheless, filter-artefact-caused spurious CFC, in particular CFS, is difficult to dissociate from genuine cross-frequency coupling by inspection of the waveform shape of any single signal in isolation. Local CFC estimates are thus prone to ambiguous results. However, CFC is necessarily genuine when there is evidence for two distinct coupled processes. Building on this notion, we advance here a conservative test to identify genuine CFC, *i.e.*, one that minimizes false positives, based on connection-by-connection testing of whether CFC can unambiguously be attributed to two separable processes. In this study, our objectives were (i) to investigate if genuine inter-areal CFC between brain regions characterizes in meso- and macro-scale resting-state activity in human SEEG and source-reconstructed MEG, respectively, (ii) to reveal the functional organization of these networks, (iii) to test whether the two predominant forms of CFC, PAC and CFS, were phenomenologically similar, and (iv) to investigate whether the strength of resting-state CFC is predictive of cognitive performance.

We estimated whole-brain connectomes of CFS and PAC and identified anatomical and topological structures therein. With SEEG data, we further addressed the putative distinct roles of generators in deep and superficial layers. We found that genuine inter-areal CFS and PAC indeed characterize human resting-state activity in both MEG and SEEG after pruning of connections that could be artefact-related false positives. CFS and PAC networks were characterized by directional coupling between the prefrontal, medial, visual and sensorimotor cortices, but crucially with distinct spectral profiles and opposing directionalities. The strength of large-scale CFS RSNs was also predictive of cognitive performance in neuropsychological assessments. These results reveal the organization of genuine CFC in human resting-state brain activity and provide evidence for CFS and PAC being functionally distinct mechanisms in the coupling of neuronal oscillations across frequencies.

## Results

### A graph-theory based method for identifying genuine neuronal inter-areal CFS and PAC

Our first objective was to assess the presence and large-scale organization of genuine CFC in human resting-state brain activity at meso-scale resolution with SEEG and at macro-scale resolution with source-reconstructed MEG data. In order to systematically address the possibility that observations of CFC might be spuriously caused by filtering artefacts stemming from non-sinusoidal or non-zero-mean waveforms, we advance here a new method to control for spurious connections. Our method is based on the core assumption that any genuine CFC interaction takes place between two distinct processes, whereas spurious CFC is a property of a single process with signal components distributed to distinct frequency bands because of filter artefacts from non-sinusoidal or non-zero-mean signal properties.

Thus, we set out to test whether observations of inter-areal CFC reflect origins in two separable processes or within one process. We advance here a graph theoretical, network-motif based approach, where we assess for each observation of inter-areal CFC between areas A and B whether there is also observed inter-areal within-frequency phase synchrony (PS) and local CFC that together may lead to a spurious observation of inter-areal CFC. If this is the case, the observed inter-areal CFC possibly does not connect distinct oscillatory processes and may hence be spurious, whereas the absence of either PS or local CFC indicates that the observed inter-areal CFC cannot be attributable to a single source and is thus genuine.

Fig 1 shows basic schemata for PS, CFS and PAC (Fig 1a) and our approach for dissociating the genuine from putatively spurious observations. Spurious observations of *local* CFC occur when a non-sinusoidal signal is filtered, which creates coupled signals at distinct frequencies that cannot be easily distinguished from genuine observations of local CFC (Fig 1b). On the other hand, *inter-areal* CFC, connecting locations giving rise to separable signals, can be proven to be genuine if it can be shown that the signals unambiguously originate from separable neuronal processes (Fig 1c). Spurious inter-areal CFC may arise only when spurious local CFC is observed at one or both locations that are also inter-areally coupled via 1:1 phase synchrony (PS) either at low frequency or high frequency so that a “triangle motif” with the observed (spurious) inter-areal CFC is formed (Fig 1d-e).

We therefore developed a graph-theory based approach to identify all CFC-PS network triangle motifs that might contain spurious inter-areal CFC (Fig 1f-h). This approach only identifies those inter-areal CFC observations as genuine, which are not part of a full triangle motif (Fig 1f), whereas all others are discarded. Since this may include also genuine connections (Fig 1g) among the spurious ones (Fig 1h), our approach is conservative and provides a lower bound for the number of genuine connections. Notably, also cases wherein there is non-sinusoidal activity at both locations are excluded (Fig 1h, right half).

### Simulation of CFC and spurious interactions with Kuramoto model

We thus posit that genuine CFC may be unambiguously observed between sources that are anatomically separable because that enables the separation of the low- and high-frequency processes. The central statistical consideration in this is that observations of significant inter-areal CFC may also spuriously arise from the combination of adequately strong inter-areal 1:1 phase synchronization and local CFC, either genuine or artificial. Such spurious inter-areal CFC should always, however, be weaker than the local CFC because it arises only indirectly from statistical regularities.

To test this rigorously, we developed a generative four-population-Kuramoto model [71] for investigating the joint effects of within- and cross-frequency phase coupling (see Methods I). The model comprised of two “areas” that each contained two populations of weakly coupled oscillators; one at low frequency (LF) and another at high frequency (HF) so that *f*_HF_ = 2·*f*_LF,_ *i.e.*, with the *n*:*m* ratio (hereafter defined as LF:HF ratio) of 1:2 (Fig 2a). The populations were coupled with coupling strengths *ε* with each other via 1:1 PS, local CFS and inter-areal CFS, and these couplings varied with a shared coupling factor *c*. The model produced salient 1:1 and 1:2 phase coupling at large coupling values (all *ε* = 0.5, *c* = 0.3) with biologically plausible intermittent synchronization dynamics (Fig 2a, right).

**Fig 2.**
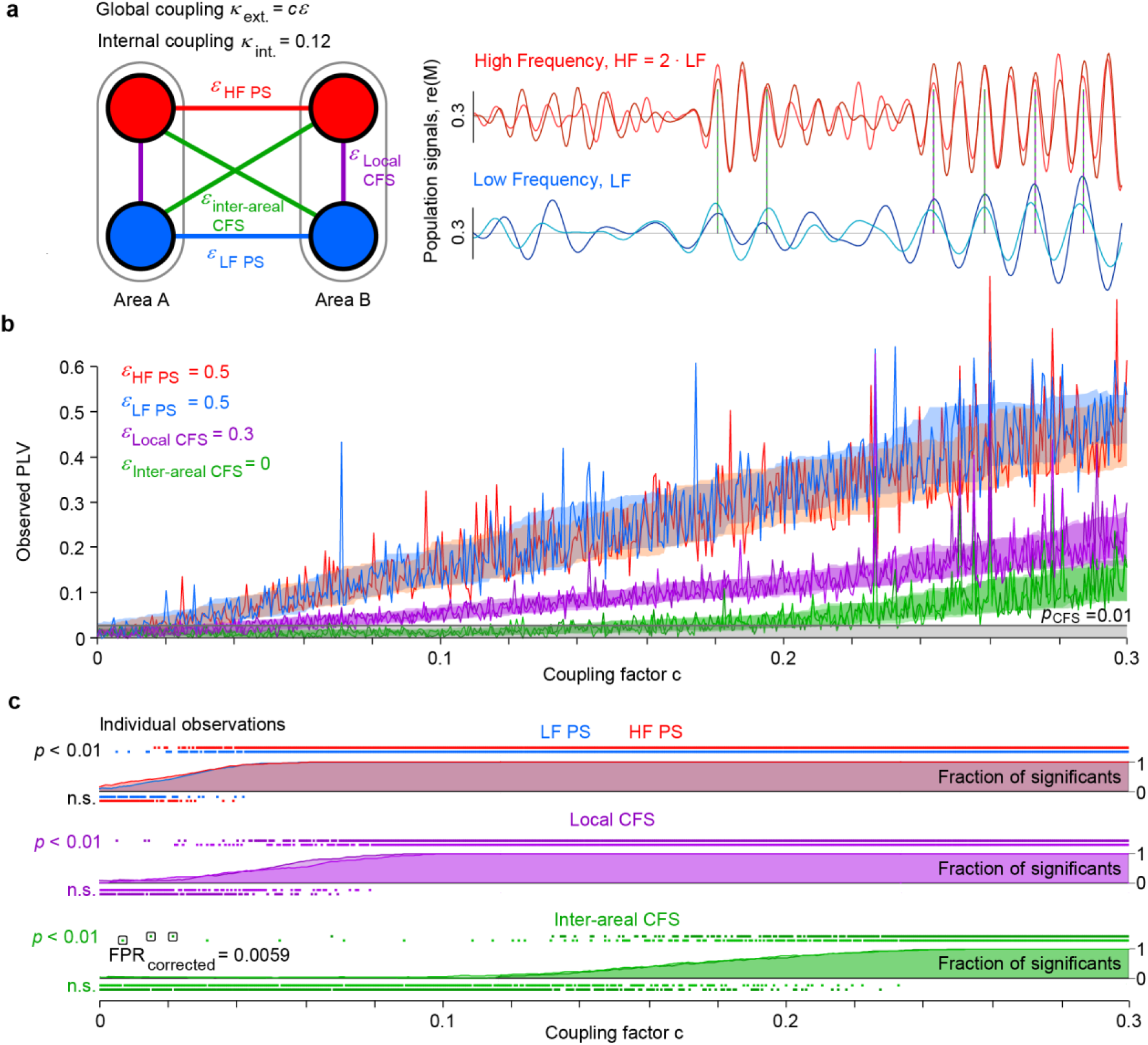
Generative modelling of joint 1:1 and 1:2 cross-frequency phase coupling. **a)** Two areas (A and B), each containing two populations of Kuramoto oscillators (*N* = 500) at low and high frequencies: LF, HF. The populations exhibit intermediate and intermittent internal synchronization (see time series) and are mutually coupled by population signal based 1:1 PS or 1:2 CFS phase coupling (see *ε*). **b)** Increasing the 1:1 and local CFS coupling between populations led to strengthening of the corresponding phase correlations (PLV, red and blue lines for PS and purple for local CFS) and, in the regime of strong coupling, also to the emergence of spurious inter-areal CFS (green lines). Each data point indicates the observed phase correlation (PLV) in a single simulation with 100000 iterations (5000 cycles of the fast oscillation) with random initial parameters in a series of 512 simulations for coupling factors from 0 to 0.3. Shaded areas indicate the 16th and 84th PLV percentiles across simulations. The gray line and shaded area indicate the PLV threshold for nominally significant CFS at *p* < 0.01. **c)** Phase correlation statistics: small squares indicate significant (*p* < 0.01) or not-significant (*n.s.*) phase correlation observations in individual simulations in the example of series of panel b. Lines and the shaded areas indicate the fraction of significant observations as a function of the shared coupling factor *c*. Black frames indicate the observations of inter-areal CFS that were not associated with significant local CFS and 1:1 PS and would thus remain as false positives after the correction proposed in this study.

To investigate the emergence of spurious inter-areal CFS, we simulated inter-areal PS and local 1:2 CFS with zero genuine inter-areal CFS. Hence here all observations of inter-areal CFS were spurious and driven by the indirect joint effect of local CFS and inter-areal PS (Fig 2b). This analysis showed that as 1:1 PS and local 1:2 CFS increased (see Fig. 2b, *c* > 0.02), also inter-areal 1:2 CFS was indeed observed in increasing strength. The crucial test for our method was to then inspect the significant (nominal *p* < 0.01) observations of inter-areal CFS. For each such observation, we tested whether it would be excluded by a simultaneous observation of significant local CFS and significant inter-areal PS. We found that this was, by and large, the case (Fig 2c). Because inter-areal CFS arose through indirect effects of local CFS and inter-areal PS that reached significance at much lower coupling values, essentially all spurious inter-areal CFS observations were correctly rejected (example false positives encircled in Fig. 2c bottom panel). The nominal false positive rate was 0.007 ± 0.002 (mean ± SD) across the simulations, and these false positives were attributable to the very-low coupling regime where both local and inter-areal CFS were at chance level. Hence for couplings above chance level, the method proposed here is effective in pruning putative spurious observations of inter-areal CFS.

### Inter-areal CFC in single-subject MEG and SEEG analyses

To estimate the presence of genuine inter-areal CFC and to map the functional organization CFC networks, we used eyes-open resting-state MEG data (10 min, 19 subjects) from healthy controls and eyes-closed resting-state SEEG data (10 min, 59 subjects) from epileptic patients (see S1 Fig and S2 Fig for the analysis workflow). SEEG data analysis was performed at the level of individually localized electrode contacts from which we excluded those located in the epileptic zone or exhibiting large artefacts (see Methods II–III). For the MEG subjects, also MRI scans were obtained and used for generating individual cortical source models for source reconstruction (see Methods IV). We obtained for each subject a cortical parcellation (see Methods V) of 200 cortical parcels by iteratively splitting [72] the largest parcels in the Destrieux atlas [73].

For both SEEG and MEG, the broadband time series were filtered into narrow frequency bands from 1 to 315 Hz. For MEG data, these were then inverse-modeled and collapsed to cortical-parcel time series (Methods VI– VIII). We excluded from further analyses parcels and parcel-parcel connections for which the source-reconstruction and connection-detection accuracy was poor, respectively (see Methods IX). From these data, we estimated inter-areal CFC between all remaining SEEG electrode contacts and parcels of MEG data. CFC was estimated for low-frequency (LF) time series between 1–95 Hz and high-frequency (HF) time series between 2–315 Hz at LF:HF ratios of 1:2–1:7. For the removal of the spurious connections as described above, we also estimated inter-areal 1:1 phase synchrony (PS) and amplitude envelope correlations (AC) between pairs of electrode contacts or parcels (see Methods X-XII).

To first acquire proof-of-concept for genuine CFC at single-subject level, we identified single-subject datasets with strong CFC. We selected a MEG participant with strong inter-areal CFS between alpha (α) and beta (β) oscillations with a ratio of 1:2 (α:β CFS) and an SEEG participant with strong inter-areal PAC between α and gamma (γ) oscillations (α:γ PAC) with ratio of 1:5. We focused on observations of inter-areal CFC between areas that were *not* connected by inter-areal PS and/or local CF in the triangle motif, so that CFC between them was genuine from the perspective of our approach (see Methods XIII). To then measure CFS in an independent manner that allows the dissection of filter artefacts from genuine coupling, we averaged unfiltered data segments time-locked to the peaks of narrow-band filtered α oscillations in the temporal sulcus (TS) (see Methods XIV). Time-frequency (TF) analysis of the average signal revealed a peak only in the α band, showing that neither genuine nor spurious local CFS was observable therein (Fig 3a). We next used the same α oscillation peak latencies to average broad-band signals from another source, central sulcus (CS, Fig 3b). TF analysis of the peak-averaged broad-band data in CS revealed an oscillation in the β-band, matching averaged β-band filtered data. However, no components in the α band in CS were found in time-frequency analysis, which confirmed the absence of both local CFS therein and inter-areal α PS between TS and CS. The observation of α-peak-locking of β oscillations between these two regions thus unambiguously indicated genuine inter-areal CFS coupling. As a confirmatory analysis, we estimated time-resolved α:β CFS between these two locations and found α:β CFS to be significant at *p* < 0.01 for a duration of ∼300 ms around the α peak in TS (Fig 3c).

**Fig 3.**
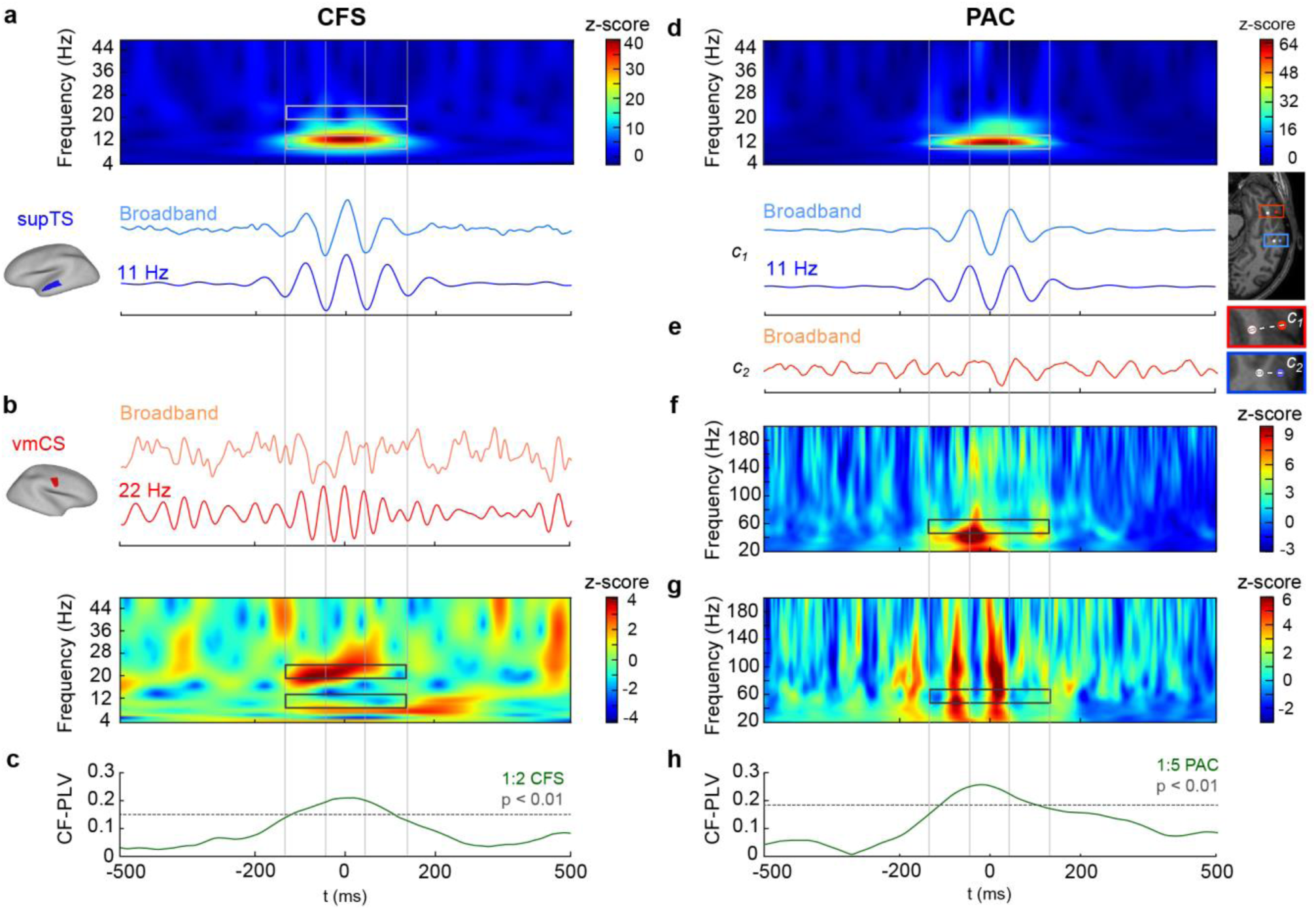
Genuine inter-areal CFS and PAC at the single-subject level. **a)** Averaged broadband time series and α**-**band (11 Hz) filtered LF time series time-locked to the α peaks in the left superior temporal sulcus (supTS) for a representative MEG subject. Time-frequency representation (TFR) of the broadband average reveals α oscillations (highlighted by lower grey box), but an absence of β oscillations (in area of upper grey box), and thus the absence of both non-sinusoidal filter artefacts and genuine local 1:2 CFS. **b)** Averaged broadband-filtered and β-band-(22 Hz) filtered HF time series for right ventromedial central sulcus (vmCS) time-locked to the α LF peaks identified in supTS. TFR of the broad-band average reveals β oscillations but an absence of α oscillations. Thus, vmCS shows no filter artefacts, local CFS, or inter-areal α or β CFS (highlighted by grey boxes). **c)** PLV time-series (green line) for 1:2 α:β CFS for the LF and HF time series averaged over α-peak time-locked segments. Dotted gray line shows the PLV value above which CFS is significant at *p* < 0.01. **d)** Averaged broadband filtered and α**-**band (11 Hz) filtered LF time series time-locked to the α troughs in an electrode contact *c*_*1*_ located in the middle temporal gyrus (mTG) in a representative SEEG subject. The electrode location is marked by the red circle on the brain, and the closest white matter contact (used as reference) by the white circle next to it. TFR of the averaged broadband time series reveals α oscillations (highlighted by grey box), and an absence of higher frequency components suggesting the absence of systematic non-sinusoidal filter artefacts that would show up as local CFS here. **e)** Averaged broadband time-series in an electrode contact c_2_ in inferior temporal sulcus (iTS) that is time-locked to the α troughs in contact c_1_ reveals no clear α oscillations, showing an absence of α PS between *c*_*1*_ and *c*_*2*_. The location of electrode contact c_2_ is marked by the blue circle on the brain, and the closest white matter contact (reference) by the white circle next to it. **f)** TFR of oscillation amplitudes in *c*_*1*_ time-locked to α peaks shows little modulation of γ amplitudes by α phase at frequencies above 40 Hz. **g**) TFR of oscillation amplitudes in *c*_*2*_ show co-modulation of γ amplitudes in *c*_*2*_ and α cycles (*i.e.* α phase) in *c*_*1*._ The frequency region at around 55 Hz, where the HF of the 1:5 PAC should be seen, is marked with grey boxes. **h)** PLV time series (green line) for 1:5 PAC between LF time series in *c*_*1*_ and LF-filtered amplitude envelope of 55 Hz NB in *c*_*2*_. Dotted gray line shows the PLV value above which PAC is significant at *p* < 0.01. Plot data is available online [115].

We then adopted this approach to assess local and inter-areal PAC. We first detected α troughs from SEEG data and averaged data segments time-locked to these troughs in the electrode contact *c*_*1*_ located in the middle temporal gyrus (mTG). Both the averaged broadband time series itself and its TF analysis showed only α oscillations (Fig 3d). The broad-band signal in an electrode contact *c*_*2*_ in the inferior temporal sulcus (iTS), time-averaged to the α troughs identified in the first contact, revealed no salient α oscillations, showing that these two contacts were not coupled by α PS (Fig 3e). Amplitude TF analysis, *i.e.*, averaging of the narrow-band amplitude envelopes around the α peaks in *c*_*1*_, revealed no evidence of local PAC in *c*_*1*_ (Fig 3f) save for a peak at around 40 Hz during one α cycle. However, in *c*_*2*_, γ amplitude was co-modulated by multiple *c*_*1*_ α cycles over a wide range of frequencies, including 55 Hz (which was the initial finding), indicating the presence of true inter-areal PAC (Fig 3g). As a confirmatory analysis, then evaluated time-resolved 1:5 α:γ PAC between LF 11 Hz at *c*_*1*_ and HF 55 Hz at *c*_*2*_ (frequencies indicated by the grey boxes in Fig 3d and Fig 3f,g respectively) and found that PAC was significant for nearly 3 α cycles at *p* < 0.01 around the central α trough (Fig 3h).

### Connectomes of inter-areal CFS and PAC in SEEG and MEG data

With both theoretical support for our method and experimental proof-of-concept for CFS and PAC, we mapped the CFC connectomes (*i.e.*, CFS and PAC between all parcels / channels) for each subject, for all low frequencies (LF) between 1–95 Hz and for all LF:HF frequency ratios 1:2–1:7, in both SEEG (*N* = 59) and MEG (*N* = 27) data. In order to first quantify the prevalence of significant CFS and PAC connections in the SEEG and MEG data, we compared the CFS and PAC connectomes against individually-generated surrogate data and identified statistically significant (*p* < 0.01) connections separately for each subject (see Methods XV). We denoted the proportion of significant CFS and PAC connections from all possible connections as the connection density, *K*. To represent these data at the group level, we averaged the individual *K* values and plotted group-level connection density spectra (*K)* as function of LF separately for each LF:HF ratio (Fig 4, shaded areas indicate 95% confidence limits of the mean estimated by bootstrap resampling). *K* spectra thus summarize the group-mean *extent* of significant CFC in individual cortical networks.

**Fig 4.**
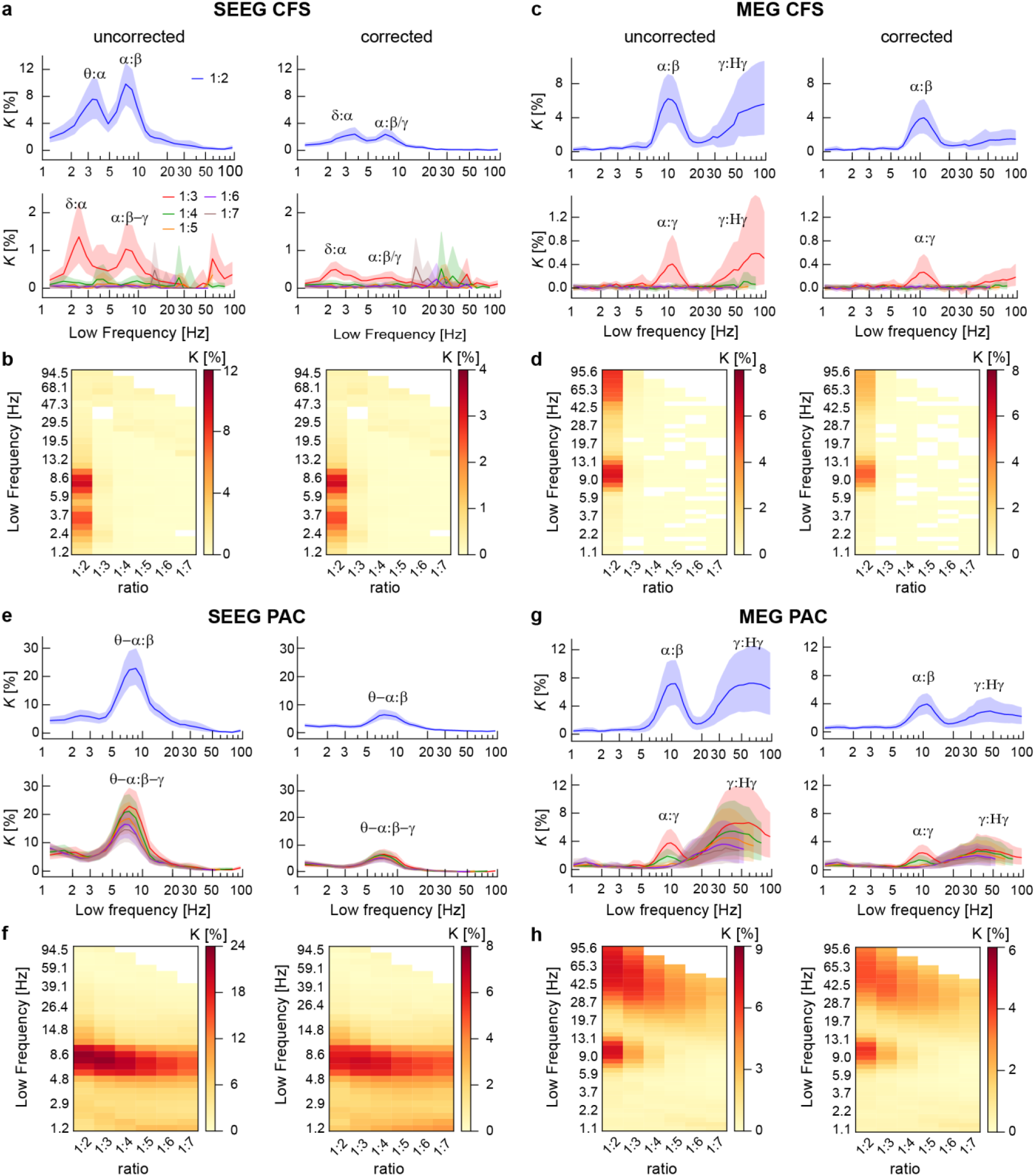
Genuine observations of inter-areal CFS and PAC. **a)** Connection density (*K*), *i.e.*, the fraction of significant connections over all possible connections, of inter-areal CFS in SEEG at the group-level before (left) and after removing possible spurious connections (right) for LF:HF ratio 1:2 (top row) and ratios 1:2–1:7 (bottom row). The *x*-axis shows the low frequency (LF). *K* values are plotted with 95% confidence limits obtained from surrogate data. **b)** The same data as in A) but presented in a matrix where each frequency-ratio combination is a matrix element. *K* is again presented before (left) and after removal of possibly spurious connections (right). **c-d)** Inter-areal CFS in MEG before and after removing possibly spurious connections. Robust α:β CFS at ratio of 1:2 and α:γ CFS at ratios 1:3 characterize SEEG and MEG data before removing spurious connections. Although *K* is reduced by removing the putatively spurious connections, α:β at 1:2 ratio and α:γ CFS at 1:3 ratio remain significantly above zero. **e**-**f)** Inter-areal PAC in SEEG data and **g**-**h)** in MEG data before and after removing spurious connections as in a-d. SEEG is characterized by robust PAC of θ–α oscillations to higher frequencies in α–γ bands at ratios 1:2–1:7, indicating that α–γ band amplitudes are modulated by phases of θ–α oscillations. In MEG, PAC is observed between α phase and β–γ band amplitudes at ratios 1:2–1:4, as well as between γ and Hγ oscillations at all ratios. The connection densities are reduced by removing putatively spurious connections but remain significantly above the zero. Plot data and underlying connectome data are available online [115].

In both SEEG (Fig 4a,b) and MEG (Fig 4c,d), the connection density spectra revealed a LF α peak at ratios 1:2 and 1:3, which indicates significant CFS coupling between α with β and γ oscillations i.e., α:β and α:γ CFS. In SEEG, the frequency range of this peak was approximately 6–12 Hz and in MEG, 7–15 Hz. This elevated amount of significant inter-areal α:β and α:γ CFS was also reflected as a peak in the corresponding graph strength (*GS*), which shows that these CFS connections were also stronger in terms of the phase-locking strength (S3 Fig). We also observed similar α-band peaks in *K* spectra of inter-areal within-frequency PS (S4 Fig) and in estimates of local CFS (S5 Fig). In addition to α-oscillation based CFS, we found in SEEG LF peaks in the range 2–5 Hz, covering parts of both delta (δ) and theta (θ) bands (Fig 4a,b). These δ–θ oscillations were synchronized at ratios 1:2 and 1:3 with θ–α oscillations, although no δ–θ peak was found in the within-frequency PS analysis (S3 Fig). In MEG, on the other hand, CFS was significant also among γ and high-γ bands at ratios 1:2 and 1:3, although the wide confidence limits indicated large inter-individual variability (Fig 4c,d). We next assessed inter-areal phase-amplitude coupling (PAC) with the same approach as above and found significant PAC in both SEEG (Fig 4e,f) and MEG (Fig 4g,h). As the most salient peak in the connection density spectra, we found that PAC coupled the phase of θ–α band oscillations (5–12 Hz) with the amplitude envelopes of oscillations at higher frequencies. PAC in SEEG was robust throughout the studied range of LF:HF ratios and coupled θ–α oscillation phases with the amplitude of neuronal activity up to the Hγ band. In MEG, α oscillation phases (7–12 Hz) were coupled with β and γ amplitudes at ratios 1:2–1:4. In MEG, PAC also coupled γ and Hγ band oscillations, similarly to CFS. On the other hand, the δ–θ band oscillations that were coupled via CFS with α oscillations in SEEG, were not observed to exhibit PAC with these or higher frequencies, indicating that the observed δ–θ:α coupling in SEEG was specific to CFS. Overall, these data suggest that robust inter-areal cross-frequency coupling, both CFS and PAC, of θ and α oscillations with oscillations in β and γ frequencies is characteristic to human resting-state brain activity.

### Genuine inter-areal CFS and PAC in resting-state brain activity

We addressed then whether the findings of CFC were attributable to filtering artefacts or reflected genuine neuronal coupling. To remove all potentially spurious connections of CFS, we discarded observations of inter-areal CFS between such sources that were also connected by both inter-areal 1:1 PS and local CFC by using the triangle motif analysis as described above (Fig 1, see also Methods XIII). We found that after the pruning of all putatively spurious connections, the mean connection density of CFS and its 95%-ile confidence limit remained above zero for CFS for α:β CFS in MEG and SEEG and for δ:α CFS in SEEG (Fig 4a,b). In SEEG, the correction removed a larger fraction of CFS connections than in MEG, indicating that the larger initial connection density *K* in SEEG may have been due to putative spurious couplings. In MEG, observations of CFS between γ and Hγ were reduced to near zero, suggesting that this phenomenon was mostly spurious and possibly arising from muscle artefacts [74]. In summary, genuine CFS was observed in human resting state brain activity also after application of our correction method, which due to its conservative nature likely underestimates the actual number of significant genuine connections.

We next applied the triangle-motif based correction to remove ambiguous PAC connections, with the difference that amplitude envelopes (S6 Fig) were used instead of PS to detect HF interactions and PAC for detecting local CFC (see Methods XV). After applying the correction method and removing the possibly spurious PAC connections, the connection density for PAC remained significantly above zero (as indicated by the 95% confidence limits) for PAC between θ–α and α–γ oscillations in SEEG as well as for PAC between α and β–γ oscillations in both SEEG in MEG. The connection density values for PAC between γ and Hγ in MEG remained significantly above zero after removing the possibly spurious connections, although they too were greatly attenuated.

As our correction method for spurious interactions is based on estimation of PS, it is affected by the metric of PS used. For the results presented so far, we used the weighted phase lag index (wPLI) [75] that yields PS estimates which are not inflated by volume conduction (SEEG) and source leakage (MEG). However, it is insensitive to genuine zero-lag neuronal couplings, and may thus underestimate the genuine extent of PS. To test whether this is a significant confounder, we also used the phase-locking value (PLV) to estimate PS. PLV is not markedly sensitive to variation in phase difference, but it is inflated by linear mixing [76]. To compensate for this and reduce the effects of linear mixing, we excluded the signal-leakage-dominated short-range connections from analyses of MEG data (see Methods IX). Even so, we found a greater connection density for PS measured with PLV than with wPLI in MEG (S4 Fig). Correspondingly, the correction led to a greater reduction of *K* in MEG CFS (S7 Fig) but, importantly, the connection density of α:β CFS remained significantly above zero. In SEEG, the corrected *K* values for PS were more similar between PLV and wPLI, in line with the fact that in appropriately referenced SEEG, volume conduction is well controlled [6]. We also computed corrected PAC values using PLV as PS metric. Results were largely similar when PLV instead of wPLI was used for estimating LF PS (S7 Fig c,d). Taken together, these results show that using our novel method for removing potentially spurious inter-areal CFC, genuine inter-areal CFS and PAC between separable sources can be observed in both SEEG and MEG data and that our method is not qualitatively confounded by the method used for estimating within-frequency PS.

### Resting-state CFS in eyes-open and -closed conditions

In the SEEG resting state dataset used here, participants had their eyes closed to limit the disturbances typical to the clinical environment, whereas eyes-open resting state data was acquired from MEG participants for compatibility with visual tasks. As the amplitude of local α oscillations is greater in the eyes-closed than in eyes-open state [14, 18, 77], we asked whether the larger *K* values found in SEEG compared to MEG could be explained by differences in the brain state. We recorded new MEG eyes-open and eyes-closed resting state data from 10 healthy subjects and computed inter-areal CFS in the same manner as described above. Significant, and qualitatively identical, 1:2 CFS between α and β oscillations was observed in both eyes-open and eyes-closed resting state but with greater *K* values in the eyes-closed than in eyes-open condition (S8 Fig). This parallels the overall larger *K* values in SEEG data and overall shows that resting-state CFS is qualitatively unaffected by the resting-state condition. As no θ:α CFS coupling was observed in eyes-closed MEG, the θ:α CFS observed in eyes-closed SEEG does not result from the lack of visual input but is a genuine property of the meso-scale brain dynamics observed with SEEG.

### Inter-areal CFC decreases as a function of distance

Observations of significant CFC after removal of putative false positive CFC using a method that minimizes false positives strongly suggests that genuine inter-areal CFC characterizes human resting-state brain dynamics. We next set out to investigate whether CFC would be dependent on the distance between the cortical sources. Since prior studies [6, 31, 39, 78] have shown that within-frequency PS is negatively correlated with distance, we expected similar results for CFC. We divided the electrode contact pairs in SEEG and parcel pairs in MEG into three distance bins containing equal numbers of connections each, and computed 1:2 and 1:3 CFS and PAC, as well as PS, in each of these bins (see Methods XVI). CFS and PAC were observed in all distance bins in both SEEG and MEG data (Fig 5). For nearly all low frequencies, the greatest *K* values for CFC were found for the shortest distances (blue lines) and the smallest *K* values for the longest distances (green lines) both before and after removing possibly spurious connections, and all three bins were found to be significantly different pairwise from each other (Wilcoxon test, *p* < 0.05, corrected for multiple comparisons) for 1:2 CFS and 1:2–1:3 PAC. PS graph strength (*GS*) also was found to decrease with distance (S3 Fig) as observed before [6, 31, 39, 78].

**Fig 5.**
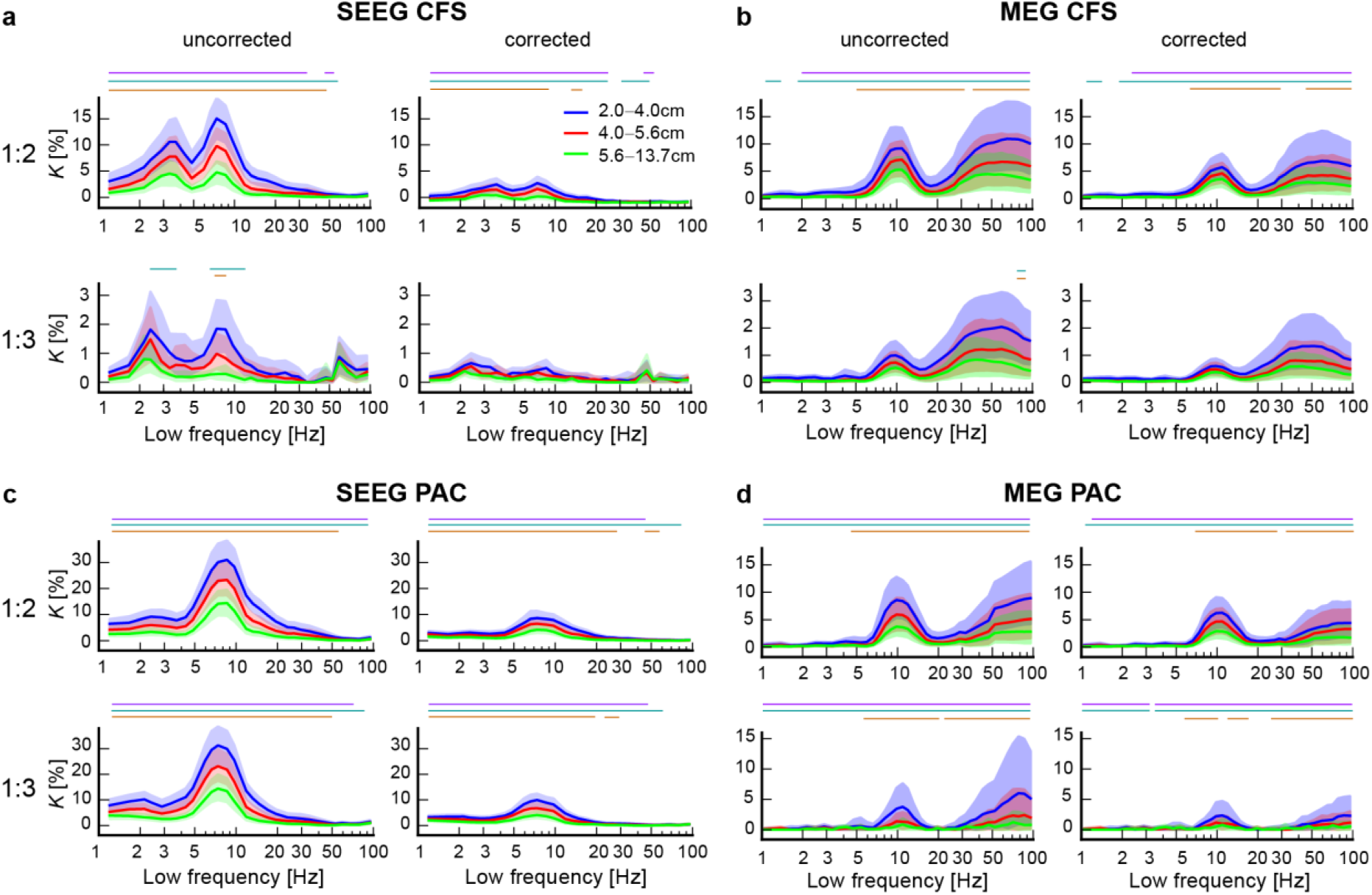
Inter-areal CFS and PAC decrease as a function of distance. **a)** Connection density (*K*) for uncorrected (left) and corrected (right) inter-areal CFS estimated separately in three distance bins containing equal numbers of connections for 1:2 (top row) and 1:3 (bottom row) inter-areal CFS in SEEG data. All values are plotted with 95% confidence limits indicated by shades. The colored bars and stars indicate LFs where *K* values between distance bins were significantly different (Wilcoxon test, *p* < 0.05, corrected) between distance bins (purple: short vs. medium; turquoise: short vs. long; orange: medium vs. long). **b)** Same as a) for CFS in MEG. **c)** Same as a) for inter-areal PAC in SEEG and **d)** in MEG. Connection density of CFC was larger for the shortest than for the longest distance bin for CFS at ratio 1:2 and for PAC at ratios 1:2 and 1:3 in both SEEG and in MEG data across most of the frequency spectrum. Plot data and underlying connectome data are available online [115].

### Inter-areal CFC is dependent on laminar depth

We then investigated whether inter-areal 1:2 and 1:3 CFS and PAC in SEEG would vary along cortical depth, which would yield insight into the underlying cortical current generators of human CFC interactions. The electrode segmentation algorithm used here enables the separation of electrode contacts in deep and superficial cortical layers (see Methods XVII), but not localization to specific layers because of the electrode size and limits of localization accuracy. Using this approach, a previous study has identified cortical-depth dependent coupling profiles for within-frequency PS [6].

We found that before the pruning of spurious connections, CFS and PAC of between δ–θ and θ–α LF oscillations with higher frequencies showed largest *K* values between the electrodes in superficial cortical layers (Fig 6, red lines) and lowest *K* values between those in the deeper layers (blue lines), this difference in *K* between being significant for θ:α and α:β CFS at ratio 1:2 and for PAC at 1:2–1:3 over a wide LF range (Wilcoxon test, *p* < 0.05, corrected for multiple comparisons). Values for CFS connections between electrodes in more superficial and deeper layers (green and purple lines) laid in between these values. After the pruning of possibly spurious connections, however, these differences were less pronounced and did not exceed the significance threshold. For PAC, connections with the LF electrode in a deeper and the HF electrode in a more superficial layer were now most prominent and those with the inverse relationship least prominent, which was significant over α:β PAC. Thus, while CFC was indeed dependent on cortical depth, further studies are needed to clarify their source in the cortical laminae with more precision as well as their dependence on inter-areal PS and local CFC in the correction approach.

**Fig 6.**
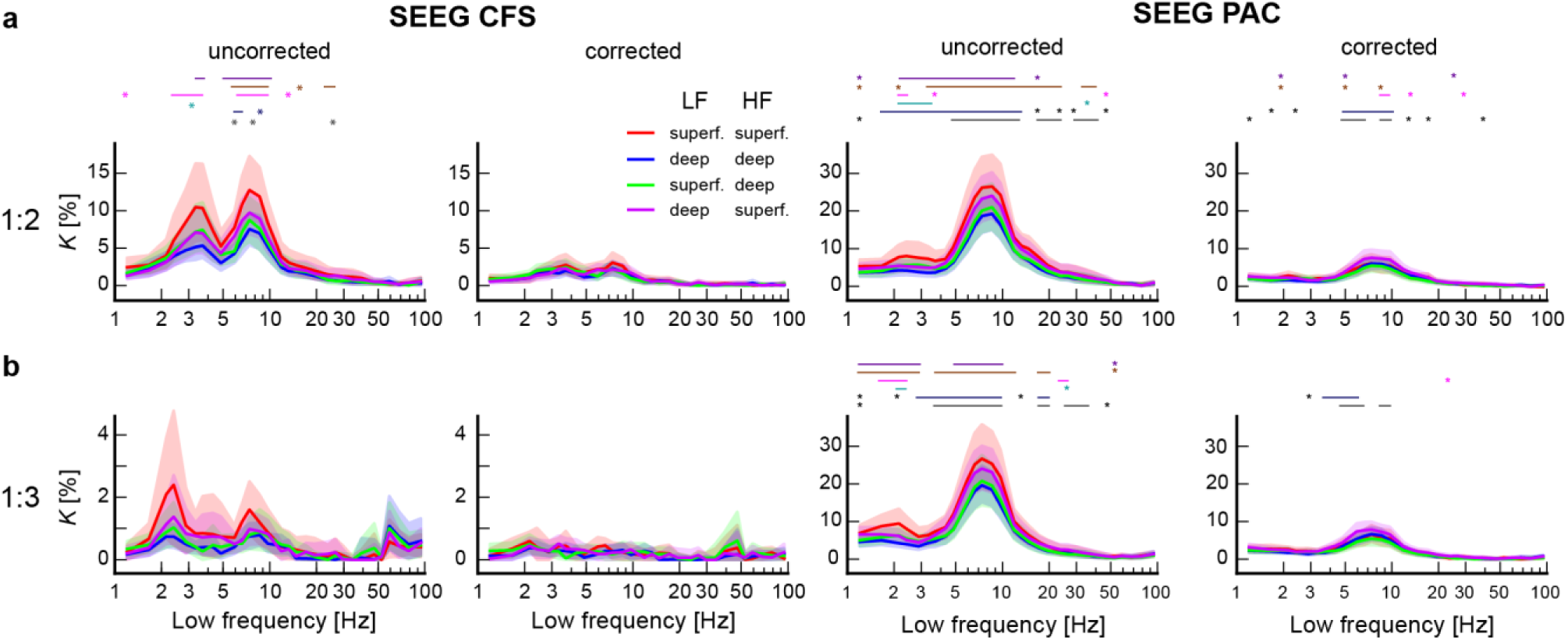
CFS and PAC in different laminar depths in SEEG data. **a)** Connection density *K* for uncorrected (left) and corrected (right) inter-areal CFS at ratio 1:2 (top row) and 1:3 (bottom row) in SEEG among electrode pairs that were either both in more superficial (s) layers (red), both in deeper (d) layers (blue), where the LF electrode was in a more superficial layer and the HF electrode in a deeper layer (green) and vice versa (purple). The colored bars and stars indicate LFs where *K* values were significantly different between laminar depth combinations (Wilcoxon test, *p* < 0.05, corrected) (purple: s-s vs. d-d; beige: s-s vs. s-d; pink: s-s vs. d-s; turquoise: d-d vs. s-d; dark blue: d-d vs. d-s; grey: s-d vs. d-s). **b)** Same for inter-areal PAC in SEEG data. In both CFS and PAC, for the uncorrected data the connection densities were highest for connections where both electrodes were localized within superficial layers and lowest for both localized within deeper layers. In corrected PAC, *K* was highest when LF electrodes were localized to deeper and HF to superficial layers and lowest when vice versa. Plot data and underlying connectome data are available online [115].

We then asked whether MEG could be preferentially sensitive to CFC interactions from either the deeper or more superficial layers. To this end, we measured with parcel degree how central each parcel was in the CFC networks and estimated the correlation between these degrees in MEG and in each of the four possible laminar depth combinations in SEEG data (Spearman’s rank correlation test) for α:β and α:γ CFS and PAC (see Methods XVIII). Parcel degree values of α:β CFS in MEG data were positively correlated with degree values when both electrodes were localized in deeper layers and negatively when they were both localized in more superficial layers (S9 Fig, *p* < 0.05, corrected with Benj.-Hochberg). The difference between the correlation values *r* between different layer combinations was determined to be significant at *z* > 1.96 with a Fisher’s z-transform for 1:2 CFS (indicated by black bar in S9 Fig). Also for α:β and α:γ PAC, the parcel degree values of MEG data were positively correlated when both electrodes were localized in deeper layers. These findings thus extend to CFC the notion that MEG may be most sensitive to neuronal current sources in deep cortical layers [79].

### Low- and high-frequency hubs differ between CFS and PAC

Finally, we aimed to elucidate the anatomical-topological organization of the δ–θ:α, α:β and α:γ CFS and PAC connectomes. We first used a conventional in- and out-degree-based graph theoretical approach [80] to estimate LF and HF centrality across the cortical surface (see Methods XIX). We represented both SEEG and MEG CFC connectomes in the 148-parcel Destrieux atlas and estimated relative directed degrees. This thus yielded, for each of the main LF peaks, what a measure of whether a given parcel was predominantly a HF or a LF hub in the CFC network. For CFS networks, HF β and γ hubs (red) were largest in somatomotor (SM) regions, posterior parietal cortex (PPC), and temporal cortex (Fig 7). LF α hubs (blue) were localized to the lateral prefrontal cortex (lPFC) and medial parietal cortex (MPC) in both SEEG and MEG, and in MEG also to the occipital cortex. Hub localization of θ:α and δ:α CFS (which was only observed in SEEG) was similar to that of α:β CFS. However, for PAC, we observed largely an opposite localization of LF and HF hubs in most cortical regions. We found the LF α hubs to be consistently localized to SM, PPC and occipito-temporal regions in both SEEG and MEG, and the HF β and γ hubs mainly to be localized to frontal regions and MPC. In order to confirm the similarity between SEEG and MEG data and the dissimilarity between CFS and PAC, we computed the correlation of relative directed degree values across parcels using Spearman’s test. A significant (*p* < 0.05) positive correlation between SEEG and MEG was found for α:β and α:β PAC, and a non-significant one for α:β CFS (Table 1). Relative degree values for α:β CFS and PAC were indeed significantly anti-correlated both in SEEG and MEG (*p* < 0.05), and also for α:γ CFS and PAC in SEEG, but not in MEG (Table 2).

**Table 1:** Parcel values are correlated between SEEG and MEG data.

|  | Rel. dir. degree |  | Directionality |  |
| --- | --- | --- | --- | --- |
|  | **r** | **p** | **r** | **p** |
| 1:2 CFS | 0.136 | 0.101 | 0.094 | 0.254 |
| 1:2 PAC | **0.233** | **0.004** | **0.259** | **0.002** |
| 1:3 CFS | -0.052 | 0.531 | 0.130 | 0.114 |
| 1:3 PAC | **0.203** | **0.013** | **0.314** | **<10<sup>-4</sup>** |

*Values obtained with Spearman's test. Significant correlations ( $p < 0.05$ ) in bold.*

**Table 2:** Parcel values are anticorrelated between CFS and PAC.

|  | Rel. dir. degree |  | Directionality |  |
| --- | --- | --- | --- | --- |
|  | **r** | **p** | **r** | **p** |
| 1:2 SEEG | **-0.254** | **0.002** | **-0.277** | **0.001** |
| 1:2 MEG | -0.114 | 0.168 | -0.106 | 0.200 |
| 1:3 SEEG | **-0.241** | **0.003** | 0.018 | 0.831 |
| 1:3 MEG | **-0.240** | **0.003** | **-0.252** | **0.002** |

*Values obtained with Spearman's test. Significant correlations ( $p < 0.05$ ) in bold.*

**Fig 7.**
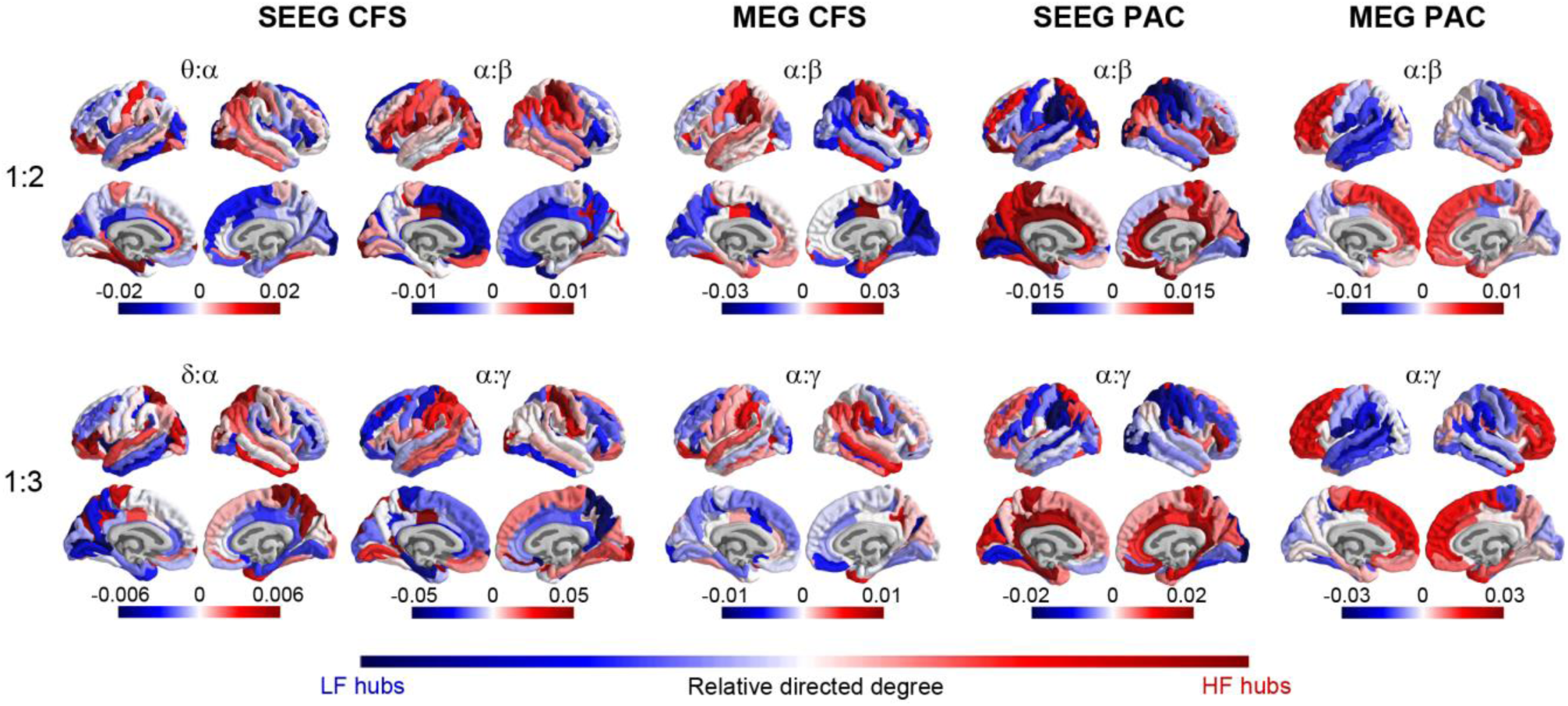
CFC networks have an asymmetric low- and high-frequency hub architecture. The functional organization of CFC networks as measured with localization of low frequency (LF) and high frequency (HF) hubs. Hubness was measured as relative LF and HF degree of each brain region (parcel). Relative degree values indicate whether a parcel is primarily a LF hub (blue) or HF hub (red) in inter-areal CFC. Top row: Brain anatomy of LF and HF hubs for CFS and PAC at ratio 1:2 connecting θ:α and α:β frequencies. Bottom row: Brain anatomy of LF and HF hubs for CFS and PAC networks at ratio 1:3 connecting δ:α and α:γ frequencies. CFS and PAC networks show saliently opposing anatomical structures connecting anterior and posterior brain regions. Plot data and underlying connectome data are available online [115].

To corroborate this graph-theoretical, parcel-degree-based approach, we then directly estimated the preferential directionality of each CFC connection between parcels and, by pooling these connections, asked whether in CFC connections, one parcel predominantly was the location of either the LF or HF oscillation. We estimated such low-vs.-high directionality for each parcel pair and each frequency pair of the main peaks across subjects (see Methods XIX). Significant directionality between the two parcels was established if the absolute directionality was higher than in 95% of permutations. We then averaged for each parcel the significant directionality values, again yielding a positive value for parcels that are predominantly HF hubs and a negative value for parcels that are predominantly LF hubs. The results were remarkably similar to those of the degree-based hubness analysis and also corroborated the salient dissociation in the directionality between CFS and PAC (S10 Fig). Estimation of similarity between SEEG and MEG, and the dissimilarity between CFS and PAC using the Spearman’s test yielded results similar to those we obtained for the directed degree (Tables 1,2). Taken together, these results provide strong evidence that the anatomy and structure of CFS and PAC connectomes are distinct.

### Resting-state CFS predict performance in neuropsychological tests

If CFC is a neuronal coupling mechanism that enables the integration of processing distributed to functionally-specialized frequency bands [9, 33-37, 39, 42, 57], its recruitment upon task demands may be limited by individual factors in a trait-like fashion. In this case, resting-state CFC could be predictive of individual performance in complex cognitive tasks that demand extensive functional integration. To investigate whether the resting-state CFC networks identified in the prior analyses were predictive of cognitive task performance, we estimated the correlation of CFS and PAC individual graph strength (*GS*) in resting-state MEG data with neuropsychological test scores collected separately (Methods XX). We computed the correlation of the test scores with CFS or PAC *GS* separately for each LF and for each frequency ratio (Spearman’s rank test). CFS between of θ–α with β–γ oscillations (θ–α:β–γ CFS) and CFS between β and γ oscillations (β:γ CFS) showed significant positive correlations with scores of Trail-Making Tests (TMT) that measure visual attention, speed of processing, and central executive functions, as well as with Zoo Map Tests that measure planning ability (*p* < 0.05, Spearman Rank correlation test, Fig 8). Intriguingly, negative correlations with the test scores were observed for CFS of α and β oscillations with higher frequencies (α–β:γ) and for γ:Hγ CFS in the Digits Tests measuring working memory performance. In contrast to CFS, PAC was largely uncorrelated with performance in any of these tests although γ:Hγ PAC was negatively correlated with performance in TMT-A (S11 Fig). For all tests together, PAC did not exceed the threshold for significance. These results suggest that in a trait-like manner, individual resting-state CFC brain dynamics are predictive of the variability in behavioral performance in separately measured tasks, which supports the notion that CFC plays a key functional role in the integration of spectrally distributed brain dynamics to support high-level cognitive functions.

**Fig 8:**
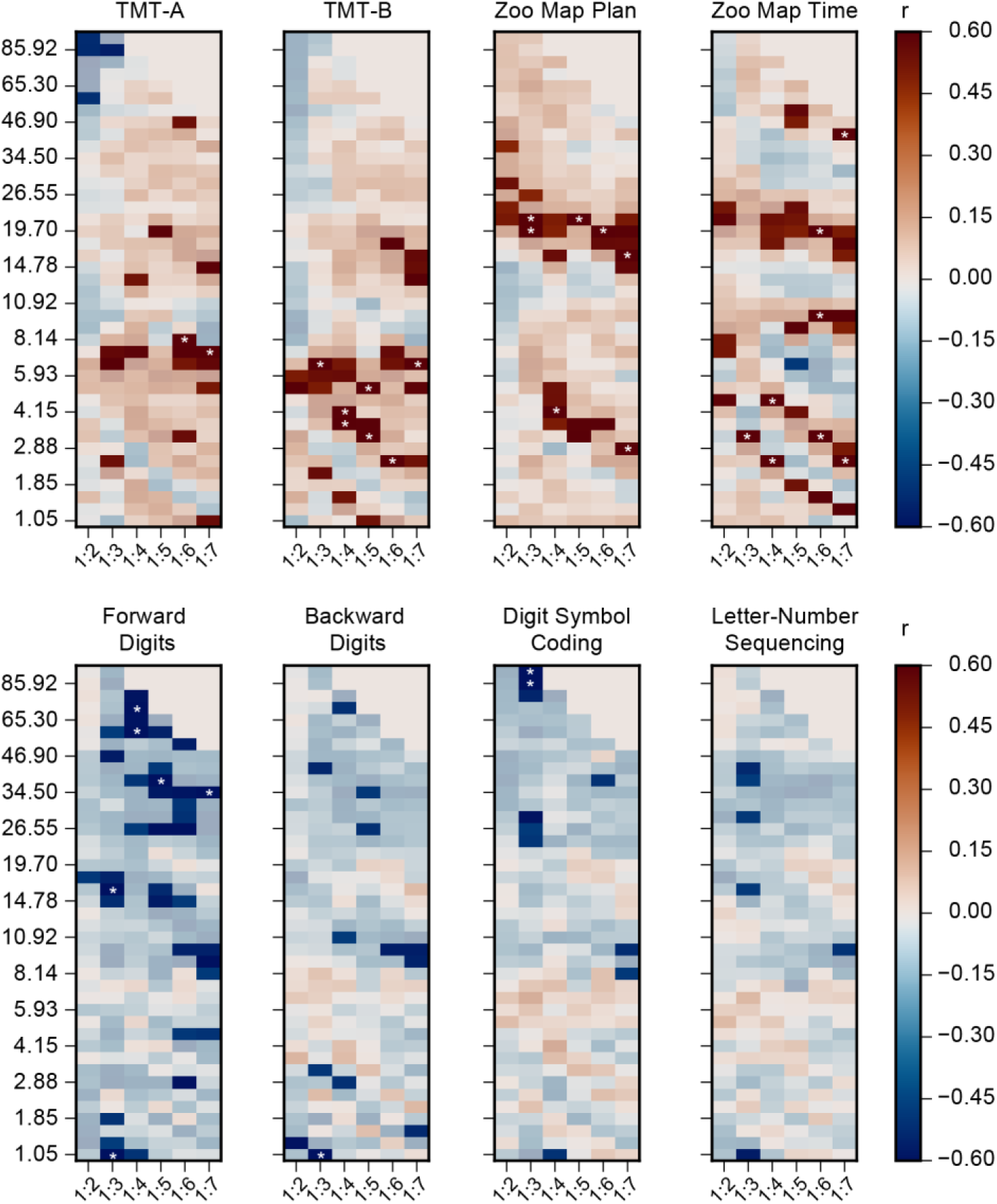
Correlation of CFS with neuropsychological test scores. The correlation of CFS graph strength (*GS*) in MEG data with scores from neuropsychological assessments for all ratios and frequency pairs (Spearman’s rank correlation test, *p* < 0.05). The assessments include Trail-Making Test (TMT) A and B, Zoo Map Plan and Time Tests, as well as Forward Digits, Backward Digits, and Digit Symbol Tests, and the Letter-Number Sequencing from Wechsler Adult Intelligence Scale–III. Red color indicates a positive correlation, so that stronger CFS is associated with better performance, while blue indicates a negative correlation between CFS and performance. Correlations with *p* > 0.05 are masked (low saturation colors). The asterisks indicate the observations that remain significant after correction for multiple comparisons (see Methods XX) across the eight neuropsychological tests and CFC frequency pairs. Plot data and underlying connectome data and neuropsychological data are available online [115].

## Discussion

Several cross-frequency coupling mechanisms, especially PAC [9, 34-37] and cross-frequency phase synchrony (CFS) [9, 38-41], have been proposed to coordinate neuronal processing across multiple frequencies and regulate the communication among coupled oscillatory networks over frequencies. Networks of phase-coupled oscillations distributed across brain areas and multiple frequencies are a core systems-level mechanism for cognition functions [11, 25-27]. Yet, only few studies have identified CFC in cortex-wide networks and across multiple frequency-pairs [39, 41, 56, 60].

We propose that that cortex-wide CFC networks are essential for coordinating computations across many frequencies and across multiple brain regions concurrently to support complex brain dynamics and cognitive functions. We report the presence and organization of two forms of inter-areal CFC, cross-frequency synchrony (CFS) and phase-amplitude coupling (PAC), in human MEG and SEEG resting-state brain activity. Importantly, also the validity of prior all CFC findings has been questioned because filtering-artefacts arising from non-sinusoidal signals lead to spurious observations of CFC.

Here were used a novel graph theoretical network-motif-based method to distinguish genuine and putatively spurious CFC in large-scale CFC networks. Genuine inter-areal CFS and PAC both characterized human resting-state activity, but showed distinct spectral profiles, anatomical architectures, and coupling directions across distributed brain regions, which strongly suggests that they originate from distinct neurophysiological mechanisms and play unique functional roles. The strength of CFS networks was also predictive of the behavioral performance in neuropsychological assessments performed separately implying a trait-like role for individual CFS in complex cognitive tasks. Overall, these data conclusively establish the presence of two distinct types of inter-areal CFC that are in a position to underlie the coordination of neuronal processing across anatomically distributed networks in multiple oscillatory frequencies.

### Large-scale CFC networks characterize human resting-state brain activity

Human brain activity during rest is characterized by intrinsically correlated fluctuations in networks of brain regions, first identified with fMRI [28, 29]. Also in human electrophysiological measurements, PS and amplitude correlations of neuronal oscillations characterize resting-state activity in a wide range of frequencies in anatomically well-delineated structures [3-8, 81] with a modular architecture and co-localized phase and amplitude correlations [31]. It has, however, remained unknown whether resting-state activity is also characterized by cross-frequency coupling (CFC) networks, and how networks formed by CFC would be organized.

We report here the presence of genuine inter-areal PAC and CFS in large-scale resting state networks. PAC of θ and α oscillations with higher frequencies was robust throughout all investigated ratios from 1:2 up to 1:7 in SEEG data and up to 1:4 in MEG data, indicating that the phases of θ and α oscillations were coupled with the amplitudes of β, γ and Hγ oscillations. PAC has been suggested to reflect the regulation of sensory information processing achieved in higher frequencies through excitability fluctuations imposed by slower oscillations [9, 10, 34, 36, 37, 42]. These results, despite methodological differences, are similar to previous findings that have reported PAC in the rat hippocampus [46, 47], non-human primates [82, 83], and in human intracranial EEG [48-51, 60] and EEG and MEG recordings [54-56, 84-86]. In contrast to PAC, inter-areal CFS connected the phase of θ/α oscillations with the phases of β and γ oscillations only with small frequency ratios (1:2 and 1:3). CFS, by definition, reflects a stable phase-difference between coupled oscillations and thus is, by definition, associated with consistent spike-time relationships between the neuronal assemblies in the two CFS-locked frequency bands [57]. The lack of high frequency ratios in CFS is not surprising because CFS necessitates the slow oscillation to have a temporal accuracy in the sub-cycle time-scales of the fast oscillation. Hence, stable phase differences over large frequency ratios may be limited by the temporal accuracy in the cellular, synaptic, and circuit mechanisms generating the slower oscillations. However, transient CFS at larger ratios has been observed during task performance [41].

CFS and PAC similarly were observed in both MEG and SEEG data, indicating that their inter-areal CFC characterizes both the meso- (mm) and macro- (cm) scale brain dynamics. Pronounced local α oscillations have been recognized as a marker of resting-state human brain activity for several decades [18, 77]. Here we show that these α oscillations are systematically cross-frequency coupled with the faster β and γ frequency oscillations across long distances in both MEG and SEEG. CFC was present in both eyes-open and eyes-closed brain states, albeit stronger in eyes-closed data putatively contributing to larger CFC values in SEEG in addition to better signal-to-noise ratio.

### Characteristics of CFC networks in resting-state human brain activity

A central unresolved question in research aiming on emergent brain dynamics is whether networks of inter-areal CFC characterize resting human brain activity with consistent large-scale architectures. Prior studies have found phase-synchronization within frequencies in SEEG [6] and in MEG data [31, 39, 78] to decrease as a function of distance. In this study, we found that the connection density of inter-areal CFS and PAC decreased with distance between cortical parcels (MEG) or electrode contacts (SEEG), similarly to that of inter-areal PS. Intracortical recordings with laminar probes have shown that oscillations of different frequencies are generated differentially across cortical layers [87-90].

We identified the depth in cortical gray matter of each SEEG electrode [6, 91] and estimated the strength of CFC within and between cortical depths. Before the exclusion of potentially spurious connections, CFS and PAC were significantly stronger among electrode pairs that were both located in more superficial layers than between pairs both located in deeper layers. However, possibly due to decreased number of samples in corrected data, and perhaps because a larger proportion of superficial connections was putatively spurious, differences between depths were not significant for CFS after correction. For PAC, we found the greatest connection density values for connections where the slow θ–α oscillations were located in the deeper layers and the fast β oscillations in the more superficial layers, and conversely the lowest values for those where it was vice-versa. This parallels earlier studies which showed that γ synchrony is strongest in superficial layers 2–3, whereas α oscillations and synchrony are generally more pronounced in the deeper layers in both monkeys and mice, although this varies somewhat with the cortical regions [87-90].

It is important, however, to note that without a current source density analysis, for which the SEEG electrode contacts and their separation are too large, and because of complex current source geometries in cortical circuitry and the volume conduction between layers [91], our findings should be corroborated and expanded in future studies addressing the neuronal sources of CFC in different cortical layers with appropriate laminar probes. Interestingly, our data showed that CFC in MEG, in terms of the cortical node-degree structure, is most similar with that observed with SEEG when both electrode contacts were localized into the deeper layers. This result strongly suggests that the sensitivity of MEG to the post-synaptic currents in large, asymmetric, and well co-oriented neurons, *i.e.* the pyramidal neurons in cortical layers 5 and 6 that are central in thalamo-cortical loops [79, 92, 93], biases the detection of CFC with MEG towards the post-synaptic currents in these neurons.

### Distinct large-scale organization of directional network architecture of CFS and PAC

To assess the large-scale architecture of the cortical CFC networks, we used two strategies for identifying the hubs of low-frequency (LF) α and high-frequency (HF) β and γ oscillations. These hubs are the brain regions where predominantly the slower (α) or the faster oscillations (β and γ) of the CFC connection are observed. We found that the LF α and HF β and γ hubs, and their directional interactions, were asymmetrically localized between anterior and posterior brain regions. The localization of hubs was largely similar between SEEG and MEG data, which corroborates the validity of these findings. Importantly, however, we observed distinct and partially opposing localization of the LF and HF hubs for CFS and PAC. In α:β and α:γ CFS, the α LF hubs were observed in PFC and medial regions that belong to the default mode network [29] or to control and salience networks in the functional parcellation based on fMRI BOLD signal fluctuations [94-96].

This is line with many previous studies that have found α oscillations in these regions to be correlated with attentional and executive functions [14-19]. In contrast, the β and γ HF hubs were found in more posterior regions such as sensorimotor region (SM) and the occipital and temporal cortices where β and γ oscillations are often associated with sensory processing [15, 20-22]. In contrast to CFS, the α LF hubs of PAC were found in occipital, temporal, and posterior parietal cortex of show pronounced whereas the β and γ HF hubs in prefrontal and medial parietal cortex. This anatomical structure between LF and HF hubs was essentially opposite to the anatomical structure observed for CFS. These results imply that both CFS and PAC contribute to the coordination of intrinsic/task-negative and extrinsic/task-positive resting-state networks [13, 28, 97] but at least partially with opposite directional roles.

### Resting-state CFS networks predict individual cognitive variability

If CFC regulates communication between within-frequency oscillatory networks to support integration of separate computational functions of cognition [9, 33-37, 39, 42, 57], it should be correlated with the psychophysical performance in cognitive tasks. We have previously shown that α-, β- and γ-band oscillations and PS networks in these frequencies are coupled via CFS and predict performance in a visual WM task. These observations imply that CFS could coordinate the representation of sensory information achieved in gamma frequencies with the executive control achieved in α-band [41]. As within-frequency PS networks during resting-state have been shown to form a core underlying the task-state networks [8, 32], we hypothesized here that CFC resting-state networks might thus also be predictive of with cognitive performance in a trait-like manner.

We thus estimated the correlation of CFS and PAC graph strengths in MEG data with the individual variability in cognitive performance in an array of neuropsychological tests. The CFS network strength was indeed predictive of test performance. CFS of θ–α oscillations with β–γ band oscillations and CFS between β and γ oscillations were correlated positively with performance in TMT-A and TMT-B as well as with Zoo-map time test that measure the interplay of visual processing speed and central executive functions. Negative correlations were found between the strength of CFS and digit test measuring working memory performance. These results thus suggest that resting-state CFS networks are indeed predictive of individual cognitive capacities in a trait-like manner.

### CFS and PAC are distinct CFC mechanisms

Many neurophysiological models have been developed to explain how the phase of slow oscillations reflecting fluctuations in neuronal excitability can regulate the power of fast oscillations usually in the gamma-frequency band via PAC [9, 33-36, 98]. PAC, by definition, is unrelated to spike synchronization between the slow and fast oscillations *per se* and reflects either the regulation of fast neuronal processing regulated by slower excitability fluctuations or the entrainment of slower oscillations by intermittent bursts of fast oscillations. We have postulated that CFS may support different computational functions than PAC [57]. CFS is a form of phase synchrony where the stable phase difference takes place between two neuronal assemblies that oscillate with an m:n frequency ratio [38, 39, 41].

Therefore, the coupling of the phases of the faster and slower oscillations indicates, by definition, that CFS will be associated with consistent spike-time relationships between the neuronal assemblies in different frequency bands. For example, for the α:β CFS, the spikes locked to the beta oscillation would have a consistent spike-time relationship with spikes locked to the alpha oscillation in every second beta cycle. In the current study, spectral and ratio differences, distinct large-scale anatomical structures and directionalities of CFS and PAC, as well as the differential correlation of the connectomes with the scores of neuropsychological assessments provide evidence for that CFS and PAC impose distinct computational functions and likely arise via separable neurophysiological mechanisms.

This conclusion is in line with prior findings during task performance where we found inter-areal CFS and PAC to show distinct spectral profiles and CFS, but not PAC, to predict working memory performance [41]. Together these results strongly suggest that CFS and PAC are not simply different operationalizations of a shared CFC process, but rather mechanistically, phenomenologically, and functionally distinct CF coupling mechanisms.

### Genuine positive inter-areal CFC is not explained by spurious connections

The main goal of this study was to investigate if genuine inter-areal CFC characterize MEG and SEEG data during resting state. Our study was motivated by the multiple concerns that have been raised about the validity of previous observations of CFC [61-69]. To map genuine large-scale networks from human SEEG and source-reconstructed MEG data, we introduced a novel graph-theory based approach to identify and exclude all putatively spurious CFC interactions. We first verified the validity of this approach using simulations in coupled Kuramoto oscillators. We then used this rationale in a novel graph-based analytical approach to control for possibly spurious CFC connections. Since our approach may also discard a subset of genuine CFC connections, it gives a conservative lower-bound estimate of the presence of genuine neuronal inter-areal CFC.

The number of significant CFC connections was indeed reduced by the exclusion of the potentially spurious corrections, indicating that part of the commonly observed CFC in MEG and SEEG data may be caused by filter artefacts arising from non-sinusoidal signal components [61-68], or by amplitude fluctuations of non-zero-mean waveforms [69]. However, our results clearly indicate that also genuine neuronal CFC characterizes MEG and SEEG data.

Our method was based on assessing for each observation of inter-areal CFC between areas A and B whether there is also observed inter-areal within-frequency phase synchrony (PS) and local CFC that together may lead to a spurious observation of inter-areal CFC. As our method uses within-frequency PS to identify the putatively spurious connections, the number of identified potentially spurious connections depends on the PS metrics. Especially in MEG, PLV connection density values were larger than the corresponding wPLI values, as PLV is inflated by volume conduction and source leakage [10, 76, 99], whereas wPLI [75] is insensitive to all linear mixing, including also true zero-lag phase coupling. Consequently, use of PLV as PS metric led to a larger reduction of CFS in MEG than correction with wPLI, but importantly, significant genuine CFC was observed with both methods. While in this study, we focused on identifying genuine inter-areal CFC in continuous resting-state data, this approach is adaptable to analyses of event-related data and may thus be used to assess the presence of genuine inter-areal CFC during task performance.

## Conclusions

We show here that large-scale networks of genuine neuronal CFS and PAC characterize human resting-state brain activity in SEEG and MEG data. Using a new graph-theoretic approach, we eliminated observations of inter-areal CFC that could be explainable by filter artefacts. The directional organization of CFC networks showed that CFC coupled slow and fast oscillations between anterior and posterior parts of the brain, suggesting that resting-state CFC coordinates intrinsic and extrinsic processing modes. The strength of CFS networks was also predictive of cognitive performance in a separate neuropsychological assessment, which implies that individual CFS is a functionally significant, trait-like property of spontaneous brain dynamics. Salient differences in spectral patterns, functional organization, and behavioral correlates demonstrated that CFS and PAC are phenomenologically and functionally distinct, and thus likely to serve complementary computational functions.

Altogether, converging results from SEEG and MEG data provide strong evidence for the co-existence of two forms of genuine neuronal inter-areal CFC in human resting-state brain activity and reveal their large-scale network organization.

## Acknowledgements

We thank Hamed Haque, Jonni Hirvonen, Sami Karadeniz, Salla Markkinen, Santeri Rouhinen, and Jaana Simola for help with the MEG recordings and Annalisa Rubino for the help with the SEEG recordings.

## Methods

### Ethics statement

Subjects gave written informed consent for participation in research studies. The recording of SEEG, CT and MRI data in Milan was approved by the ethical committee (ID 939) of the Niguarda “Ca’’ Granda” Hospital, Milan, and was performed according to the Declaration of Helsinki.

The recording of MEG and MRI data in Helsinki was approved by the ethical committee of Helsinki University Central Hospital, Helsinki, and was performed according to the Declaration of Helsinki.

#### I. Modelling

We used a Kuramoto model [71] to investigate the direct and indirect effects of within- and cross-frequency phase coupling on phase correlations observable among neuronal populations. The model was adapted from conventional Kuramoto models so that it comprised of two “areas” that each contained two populations (*N* = 500) of oscillators; one at low frequency (LF) and another at high frequency (HF) so that their frequency ratio was 1:2. The model was defined so that for each area *k, k =* 1…4, the phase of each oscillator *h, h = 1…500*, was given by:

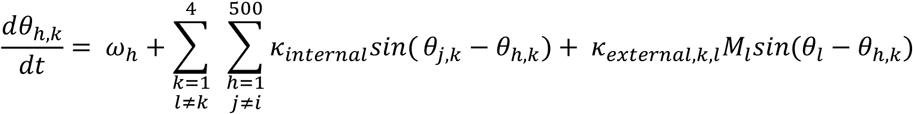

Where the phase increment per sample *ω*_*i*_ was uniformly distributed in the range from π/5*m* to π/15*m* with *m* = 1 for HF and *m* = 2 for LF. Oscillators within the populations were all-to-all connected with constant weak coupling *K*_*internal*_ = 0.12. The populations were 1:1 (*ε*_LF PS_ and *ε*_HF PS_) or 1:2 (*ε*_Local CFS_ and *ε*_Inter-areal CFS_) phase coupled with oscillators of the other populations (see Fig. 2a) so that the coupling was mediated by the population mean signals *M*_*k*_, 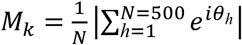. The coupling between the populations, *K*_*external*_, was varied with a shared coupling factor, *c*, so that *K*_*external*_ = *cε* for each connection specified by the corresponding *ε* value.

To validate our method for correction of spurious CFS, we set *ε*_inter-areal CFS_ = 0, *ε*_Local CFS_ = 0.3, and *ε*_LF PS_ = *ε*_HF PS_ = 0.5 in order to produce model time series that had no genuine inter-areal CFS, but rather only spurious inter-areal CFS that emerges indirectly from the combination of inter-areal PS and local CFS. We performed significance tests for PS, and both forms of CFS. We simulated 100,000 iterations of the model, yielding 5000 cycles at HF (at nominal *wi* = π/10) across 512 values of *c* from 0 to 3. Observations of PS and CFS were deemed considered significant at a nominal *p* < 0.01 obtained by setting the threshold for the observed PLV to 2.42 times the null-hypothesis PLV.

#### II. SEEG Data acquisition

Stereo-electroencephalography (SEEG) data were recorded from 59 subjects affected by drug-resistant focal epilepsy and undergoing pre-surgical clinical assessment at “Claudio Munari” Epilepsy Surgery Centre, Niguarda Hospital, Milan. Intra-cranial “monopolar” (*i.e.*, all contacts referenced to a single contact in the white matter) local field potentials (LFPs) were recorded with platinum–iridium, multi-lead electrodes with 8-15 contacts each. These contacts were 2 mm long, 0.8 mm thick, and had an inter-contact border-to-border distance of 1.5 mm (DIXI medical, Besancon, France). The neuroanatomical targets and numbers of electrodes implanted to each subject varied exclusively according to clinical requirements [100].

For each subject, one 10-minute set of eyes-closed resting state data was recorded with a 192-channel SEEG amplifier system (NIHON-KOHDEN NEUROFAX-110) at a sampling rate of 1000 Hz. The electrode contact positions were localized after the implantation by using computerized tomography (CT) scans and the SEEGA tool that performs automatic electrode contact localization and is freely available [91, 101]. Structural MRIs were recorded before implantation and rigid-body co-registration was used to co-localize MRIs and post-implant CT scans [100, 102]. Based on this, electrode contacts were assigned to one of 148 parcels of the Destrieux atlas [73].

#### III. SEEG Data preprocessing and filtering

Defective electrode contacts were identified by non-physiological activity and excluded from further analysis. For referencing, we used the closest-white matter referencing scheme [102] where each contact in cortical grey matter is referenced to the nearest contact in white matter. The seizure-onset and propagation zones were identified by clinical experts in gold-standard visual analysis and contacts in these areas were excluded from analysis, as were contacts from subcortical regions. In order to avoid spurious connectivity due to volume conduction, we excluded from connectivity analyses also contact pairs that shared the same reference or had a contact-to-contact distance < 2 cm.

We excluded all harmonics of 50 Hz line-noise using a band-stop equi-ripple finite-impulse-response (FIR) filter. Moreover, as interictal epileptic events (IIEs) such as interictal spikes are characterized by high-amplitude fast temporal dynamics as well as by widespread anatomical spread, filtering artefacts may occur around epileptic spikes and artificially inflate both PS and CFC estimates. We therefore discarded periods of containing IIEs so that we first divided the signal in non-overlapping 500 ms time windows and detected IIE events in amplitude envelopes by values that exceeded the channel mean amplitude > 5 standard deviations. Time windows where at least 10% of cortical contacts demonstrated IIE events in more than half of the 18 frequency bands were then excluded from further analyses. Time series were then filtered with Morlet wavelets with *m* = 5 using 49 roughly logarithmically spaced center frequencies from 1.2 to 315 Hz and downsampled to a sampling rate approximately 5 times greater than the wavelet center frequency.

#### IV. MEG and MRI Data acquisition

For the main study, 306 channel MEG (204 planar gradiometers and 102 magnetometers) was recorded with a Vectorview-Triux (Elekta-Neuromag) at the BioMag Laboratory, HUS Medical Imaging Center from 19 healthy participants during 10 minutes of eyes-open resting state. Overall 27 sets of resting state MEG data were obtained, with 4 participants contributing 2 sets, and 2 participants contributing 3 sets. Subjects were instructed to focus on a cross on the screen in front of them. Bipolar horizontal and vertical EOG were recorded for the detection of ocular artefacts. MEG and EOG were recorded at 1000 Hz sampling rate. T1-weighted anatomical MRI scans (MP-RAGE) were obtained for head models and cortical surface reconstruction at a resolution of 1×1×1 mm with a 1.5 Tesla MRI scanner (Siemens, Germany) at Helsinki University Central Hospital. Written informed consent was obtained from each subject prior to the experiment. In addition to the main MEG dataset, we also recorded a *de novo* 10 subject cohort (of which 4 had also participated in the main study) with 10 minute sessions of both eyes-open and eyes-closed resting-state. These data were recorded and preprocessed in a manner identical to the main MEG dataset.

#### V. Cortical parcellation

FreeSurfer software (http://surfer.nmr.mgh.harvard.edu/) was used for volumetric segmentation of MRI data, flattening, cortical parcellation, and neuroanatomical labeling with the Destrieux atlas [73]. We obtained a cortical parcellation of 200 parcels by iteratively splitting the largest parcels of the Destrieux atlas along their most elongated axis at the group-level [72, 103]. All analyses of MEG data in the main dataset were carried out using the 200-parcel parcellation, except the degree and directionality analyses (see Methods XIX) for which the data were collapsed to the original 148-parcel Destrieux atlas to facilitate the comparison with SEEG data. The analysis of the additional 10-subject MEG dataset was carried out in the 148-parcel Destrieux atlas.

#### VI. Source models and co-localization

MNE software (http://martinos.org/mne/stable/index.html) [104, 105] was used to create cortically-constrained source models, MEG-MRI co-localization, and for the preparation of the forward and inverse operators. The source models had dipole orientations fixed to pial-surface normals and a 5 mm inter-dipole separation throughout the cortex, which yielded 5086–7857 source vertices per hemisphere.

#### VII. MEG data preprocessing and filtering

Temporal signal space separation (tSSS) in the Maxfilter software (Elekta Neuromag Ltd., Finland) [106] was used to suppress extra-cranial noise from MEG sensors and to interpolate bad channels. We used independent components analysis (ICA) adapted from the Matlab toolbox Fieldtrip, http://fieldtrip.fcdonders.nl, to extract and identify components that were correlated with ocular artefacts (identified using the EOG signal), heart-beat artefacts (identified using the magnetometer signal as a reference), or muscle artefacts. After artefact exclusion, the time series data were filtered into narrow-band time series using a bank of 53 Morlet filters with wavelet width parameter *m* = 5 and approximately log-linear spacing of center frequencies ranging from 1.1 to 315 Hz. After the filtering, the time-series data were downsampled to a sampling rate around 5 times the center frequency.

#### VIII. MEG source reconstruction: inverse transform and collapsing of source signals to parcel time series

We computed noise covariance matrices (NCMs) using preprocessed (see VII) and FIR-filtered (151–249 Hz) MEG resting-state data time series. NCMs were evaluated in and averaged across 60 time windows of 10 s. This frequency band was used for NCMs because it comprises environmental, sensor, and biological noise components, but less neuronal activity than the lower frequency bands. These NCMs were then used for creating one inverse operator per subject with the MNE software and the dSPM method with regularization parameter λ = 0.11 [104, 105]. In analyses of inter-areal correlations with source-reconstructed MEG data, one confounder is posed by the spurious connections resulting from source leakage that spreads true inter-areal into false positives in their vicinity [10, 76]. In order to mitigate these effects and collapse the inverse transformed source dipole (vertex) time series into parcel time series in a manner that maximizes the source reconstruction accuracy [107], we used estimates of *vertex fidelity* to obtain fidelity-weighted inverse operators. The fidelity estimates were obtained by simulating for all 200 parcels uncorrelated, complex white noise time series (equivalent to decimated Morlet-filtered white noise), and then applying these parcel time series to all source dipoles per parcel. The source time series, *Z*_*V,orig*_, were then forward and inverse modeled (*i.e.*, considered as ground-truth parcel data, transformed into MEG sensor time series, and then source reconstructed) to obtain source time series, *Z*_*V,mod*_, that thus encompass the effects of MEG-data-acquisition related signal mixing and residual inverse-modeling source leakage. Then, *vertex fidelity* was estimated for each source vertex by the correlation between the forward-inverse-modelled data and ground-truth data. This correlation was quantified with the absolute-valued real part of the complex phase-locking value (cPLV) between *Z*_*V,orig*_ and *Z*_*V,mod*_ that was defined [38, 39, 108] as follows:

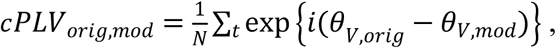

where *θ* is the phase of a complex filtered time series *Z*. Each source-dipole row of the inverse operator was then weighted with

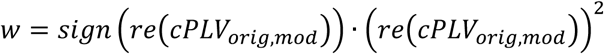

The fidelity-weighted inverse operator has higher reconstruction accuracy than a regular inverse operator for the given parcellation, because it gives greater weight to sources with better reconstruction accuracy for the signals from the parcels they belong into. The narrowband resting-state time series were inverse-modeled using this operator and collapsed into the parcellation by simply averaging the sources for each parcel. The sign operation ensures that the source current polarity switching, for example on opposing walls of a sulcus, does not result in signal cancellation at averaging.

#### IX. Removal of low-fidelity parcels and connections from MEG connectivity analysis

To focus further analyses on parcel pairs of which the interaction can be estimated with reasonable accuracy, we measured the quality of the parcellation-collapsed inverse transform with *parcel fidelity* and *cross-parcel mixing.* Similarly to the procedure for estimation of *vertex fidelity* described in VIII, we simulating uncorrelated, complex white-noise time series for all parcels, applied forward and inverse transforms, and collapsed vertex time series into parcel time series using the fidelity-weighted inverse (see section VII). We then estimated *parcel fidelity* as the absolute real part of cPLV between the original and forward-inverse modeled parcel time series for the 200 parcels concurrently, and also estimated *cross-parcel mixing* among all parcels as the |re(cPLV)| between all forward-inverse modeled time series [41].

To decrease the probability of spurious synchronization and exclude poorly source reconstructable connections, for the wPLI analysis, we excluded parcels with *parcel fidelity* < 0.1, retaining 187 of 200 parcels and 34782 (87.4 %) of all 39800 parcel pairs. For the PLV analysis, which is affected by source leakage, we additionally excluded parcel pairs with *cross-parcel mixing* of PLV > 0.2143, so that in these analyses we retained 28416 (71.4 %) of parcel pairs. The threshold for cross-patch PLV was obtained as 1.95 times the mean value in the simulations, which corresponds to a nominal *p* < 0.05 significance level. The removed parcels and connections were located mostly to deep and/or inferior sources, which are known to generate the least detectable signals in MEG and which are hence most likely to generate spurious connections [109].

#### X. Analysis of inter-areal phase-synchronization (PS)

To identify cortex-wide phase-synchrony (PS) networks, we first computed individual parcel-to-parcel (MEG data) or electrode contact-contact (SEEG data) interaction matrices. Phase synchrony (PS) was estimated using the weighted phase-lag index (wPLI) [75] and the phase-locking value (PLV) [38, 39]. Because of residual linear mixing between the parcel time series after inverse modeling, *i.e.*, source leakage, the PLV yields inflated values and artificial zero-lag false positive observations, while wPLI is insensitive to all zero-lag interactions and hence does not yield artificial interactions nor true zero-lag couplings [75, 110].

We computed PS across the whole time series for each frequency, and each contact pair *c*_*a*_, *c*_*b*_, or parcel pair *p*_a_, *p*_*b*_ with wPLI and PLV. The wPLI was defined as:

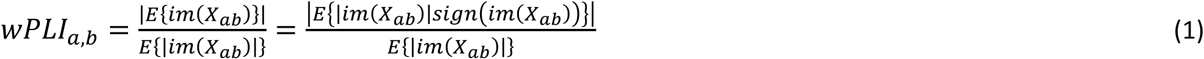

where *im* (*X*_*ab*_) is the imaginary part of the cross-spectrum of the complex time series *Z*_*a*_ and *Z*_*b*_, and *E{}* is the expectancy value operator. Here, we substituted the cross-spectrum with 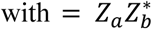, where *Z*_*a*_ and *Z*_*b*_ are Morlet-filtered narrowband time series and * denotes the complex conjugate, and used the mean over samples as the expectancy value. This can be done because Fourier- and Morlet-based spectral analysis are mathematically equivalent [111].

The PLV was defined as:

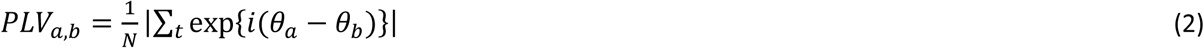

where *θ* is the phase of the complex filtered time series *Z* and *N* the number of samples. To assess the significance of synchronization at the level of individual subjects, we obtained one surrogate PS value for each contact/parcel pair where *Z*_*b*_ was randomly rotated (shifted by a random number of samples), and then calculated the means (wPLI_surr_mean_, PLV_surr_mean_) and for wPLI also standard deviations (wPLI_surr_SD_) of these surrogate PS estimates across contact/parcel pairs. Rotation, rather than more aggressive shuffling methods, was used to retain the autocorrelation structures in the data for the surrogate analyses and thereby avoid the underestimation of the null-hypothesis-level PS values. We then computed then for observed wPLI values, wPLI_obs_, the z-score as

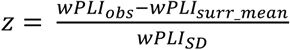

and considered those wPLI_obs_ values significant, for which z > 2, corresponding to α = 0.02. The PLV of uncorrelated phases is Rayleigh distributed and the Rayleigh distribution is only a function of its mean so that the distribution reaches the 99^th^ %-ile at PLV = 2.42 PLV_surr_mean_ [39]. We used α = 0.01 as the significance criterion for the measured PLV, PLV_meas_, and thus connections with PLV_meas_ > 2.42 PLV_surr_mean_ were considered significant. Using this approach, we obtained for each subject and each frequency the individual connection density (*K)* values, where *K* indicates the proportion of significant connections of all possible connections across channel-pairs in SEEG and parcel-pairs in MEG.

#### XI. Analysis of local and inter-areal of cross-frequency coupling (CFC): PAC and CFS

CFS and PAC were computed between all low- (LF) and high-frequency (HF) frequency pairs at ratios of *n:m* (LF:HF) from 1:2 to 1:7, and for each contact-pair *c*_*a*_, *c*_*b*_ in SEEG data and for each parcel-pair *p*_*a*_, *p*_*b*_ in MEG data. Frequency pairs were chosen so that the ratio of their center frequencies lay within 5% deviation of the desired integer 1:*m* ratio. CFS was computed as:

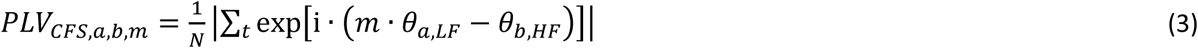

where *θ*_*a,LF*_ and *θ*_*b,HF*_ are the phases of the time series of contact /parcels. *θ*_*a,LF*_ was upsampled to match the sampling rate of the HF signal and then ‘phase-accelerated’ by multiplication with *m* [39, 41]. Local CFS (*CFS*_*loc*_) was obtained where *a* = *b* and inter-areal CFS where *a* ≠ *b*.

The strength of PAC was quantified with as:

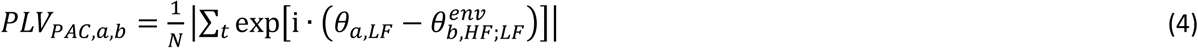

where 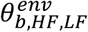 is the phase of the amplitude envelope of the HF signal filtered with a Morlet filter at LF, and downsampled to match the LF signal’s sampling rate. Local PAC was obtained where *a* = *b*, inter-areal PAC where *a* ≠ *b.*

For both CFS and PAC, we obtained, for each subject and each frequency pair, surrogate values for each contact pair or parcel pair by rotating *θ*_*b,HF*_ or 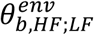 analogously to what was done for PS, and then calculated the means (PLV_CFS,surr_mean_, PLV_PAC,surr_mean_). As for PS, connections with a ratio *PLV*_*meas*_/*PLV*_*surr*_ of 2.42 or higher were identified as significant at α level 0.01 and connection density *K* was estimated as fraction of significant over possible connections, as for PS.

#### XII. Analysis of amplitude-amplitude coupling

As a pre-requisite to the removal of potentially spurious PAC (see next section), we estimated amplitude coupling (AC) with PLV as the pairwise phase synchrony of LF-modulated amplitude envelopes of the HF signals. In order to do so, we obtained the LF-filtered amplitude envelopes of the HF signals 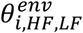 and 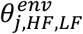, as described in the previous section, for all contact pairs or parcel pairs *i*≠*j*, and all frequency pairs LF, HF, and then computed the phase synchrony between these envelope time series using PLV. Significance for each connection, and subject-level *K* were determined as for PS.

#### XIII. Removal of potentially spurious CFC connections

The core tenet of CFC is that it indicates an interaction between two distinct neuronal processes and hence CFC is genuine when there is evidence for the presence of two separate signals. Conversely, CFC may be spurious if evidence of the presence of two separate signals is not confirmed and it remains a possibility that an observation of inter-areal CFC between two signals is due to at least one of them being non-sinusoidal and “leaking” via PS or AC to the other one. We developed a connection-by-connection test for whether inter-areal CFC can unambiguously be attributed to two separable signals. In this test, observations of inter-areal CFC are discarded between any two such signals that are also connected by both inter-areal 1:1 PS/AC and local CFC at one or both of the locations in a “triangle motif” (see Fig 1d-h). Since such a test also may remove “ambiguous” cases, where inter-areal CFC is genuine, but nevertheless part of a triangle motif (See Fig 1g), our test is conservative in the sense that it minimizes false positives, while possibly leading to false negatives, and thus provides a lower bound estimate for the number of genuine connections.

Inter-areal CFS was removed when there was a triangle motif of (significant) local CFS at the LF location and (significant) inter-areal HF PS, or when there was local CFS at the HF location and significant inter-areal LF PS (see Fig 1e-h). Likewise, inter-areal PAC was removed when there was a triangle motif of either local PAC at the HF location and low-frequency inter-areal PS, or when there was local PAC at the LF location and inter-areal HF amplitude correlation.

#### XIV. Single-subject analysis of CFC

In MEG data, we first identified a parcel pair *p*_*1*_, *p*_*2*_ with strong 1:2 α:β CFS across subjects. Next we selected a representative subject and filtered that subject’s broadband (BB) time series with Morlet filters (*m* = 5) in *p*_*1*_ at LF = 11 Hz and in *p*_*2*_ at HF = 22 Hz to obtain narrowband (NB) time series. We then identified the largest α oscillation peaks in the real-valued LF time series and averaged time-locked data segments of 1000 ms length centered around these α peaks (*N* = 62) in order to identify LF-peak-locked oscillations without a contribution from filtering artefacts.

The frequency content of the averaged BB time series at *p*_*1*_ and *p*_*2*_ was then visualized in time-frequency (TF) plots obtained by Morlet-filtering (*m* = 5) the averaged BB time series at frequencies from 4 to 48 Hz (in steps of 1 Hz) and taking the amplitude *A* of the filtered time series. Of this, we calculated, in the baseline windows from −500 ms to −200 ms and 200 ms to 500 ms relative to the peak, the mean and SD *A*_*BL_mean*_ and *A*_*BL_SD*_, and computed the z-score for *A* at each of the 1000 time points:

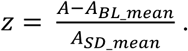

To corroborate the TF-analysis-based inferences of CFS, we also estimated the conventional 1:2 CFS (see Methods XI) with PLV between the NB time series of *p*_*1*_ and *p*_*2*_ in sample-by-sample sliding time windows of 200 ms length, *i.e.*, with phase differences concatenated across samples within the time windows and across the LF-peak-locked data segments. To estimate the null hypothesis CFS and its distribution, the above analysis was performed with randommly picked data segments of equal numbers of samples. As described in Methods X, CFS was deemed significant where PLV_meas_ > 2.42 · PLV_surr_mean_.

In SEEG data, we identified a pair of electrode contacts *c*_*1*_, *c*_*2*_ (from different electrode shafts) with strong 1:5 α:γ PAC in a single subject. We filtered that subject’s BB times series in *c*_*1*_ at LF = 11 Hz and in *c*_*2*_ at HF = 55 Hz to obtain LF (α) and HF (γ) NB time series, and in *c*_*1*_ identified the largest α troughs (*N* = 108). Centered around these troughs, we again averaged time-locked data segments of 1000 ms length and obtained a TF plot of the average of time-locked BB time series in *c*_*1*_. From both contacts’ BB time series, we further constructed BB amplitude-average TF plots by filtering the BB time series at frequencies from 20 to 200 Hz and for each of these frequencies averaging the amplitude over segments time-locked to LF (α) trough.

To again corroborate the TF analysis based inferences of PAC, we estimated 1:5 PAC conventionally between the LF NB time series in *c*_*1*_ and the averaged LF-filtered amplitude envelope of HF NB time series in *c*_*2*_ (see Methods XI) in 200 ms sliding windows, concatenated across segments as for CFS. As for CFS, we estimated the null hypothesis and deemed PAC significant where PLV_meas_ > 2.42 · PLV_surr_mean_.

#### XV. Group-level statistics

To represent these data at the group level, we averaged the individual PLV and *K* values obtained as described above in X-XII to obtain group-level graph strength *GS* and connection density *K*. In SEEG, these values were weighted with the number of contacts for each subject, which is equivalent to pooling all contacts across subjects. For estimating the group-level statistics of coupling strength and *K* for PS, AC, CFS, PAC, CFS_local_ and PAC_local_, group-level upper and lower confidence limits (2.5% and 97.5%) were computed with a bootstrapping approach, using *N* = 1000 resamplings with replacement of the subjects in the cohort separately for each LF and for each frequency-ratio LF:HF, again weighting values from SEEG with the number of contacts.

Since it can be expected that in the absence of any genuine interactions 1% of observed edges would be false positives at the significance level *p* < 0.01, we subtracted 1% from the connection density *K* of significant connections for all reported and visualized *K* values. Corrected *K* values were computed as the fraction of the remaining significant connections divided by number of possible connections after removing all putatively spurious connections using the approach described in Methods XIII.

#### XVI. Computation of CFC in distance bins

For both SEEG and MEG data, we divided all channel/parcel pairs into three distance bins of equal numbers of connections using Euclidian distance between channel or parcel pairs. For SEEG data, the distance bins were 2–4 cm, 4–5.6 cm, and 5.6–13.7 cm and for MEG data these were 0–6.3 cm, 6.3–9.1cm, and 9.1–17.7 cm. We then computed the *GS* and *K* values separately in these distance bins as described above. We tested CFS and PAC for significant differences of *K* between all 3 pairs of distance bins with a Wilcoxon signed-rank test (*p* < 0.05, corrected for multiple comparisons across frequencies and combinations with Benjamini-Hochberg) accepting the significance values if more than two consecutive LFs were significant.

#### XVII. Estimation of CFC in distinct cortical layers in SEEG data

We divided SEEG electrode contacts to two laminar depths based on the Grey Matter Proximity Index (GMPI) which is the distance between contact position and the nearest vertex of the white-grey surface, normalized by the cortical thickness in that point [91]. GMPI = 1 thus indicates the pial surface and GMPI = 0 indicates the surface between grey and white matter. Based on this depth along the cortical grey matter, contacts within 0.5 < GMPI < 1.2 were marked as “superficial” and those within –0.3 < GMPI < 0 as “deep”. We analyzed inter-areal CFC among electrode pairs in four groups: 1) where both electrode contacts were superficial; 2) where both contacts were deep; 3) where the LF contact was superficial and the HF contact deep; and 4) where the LF contact deep and the HF superficial. We tested CFS and PAC for significant difference of *K* values between all 6 pairs of laminar depth combinations with a Wilcoxon signed-rank test (*p* < 0.05, corrected for multiple comparisons across frequencies and combinations with Benjamini-Hochberg) and accepted values if more than two consecutive LFs were significant.

#### XVIII. Correlation of CFC in MEG with laminar depth in SEEG data

In order to elucidate the cortical sources contributing to the CFC connectomes observed with MEG, we performed an analysis to assess the correlation of these connectomes with those in layer-specific SEEG connectomes. We here focused on the frequency pairs that showed the most robust observations of inter-areal CFS and PAC interactions in both MEG and SEEG, namely 1:2–1:3 CFS and 1:2–1:4 PAC for the LF peak in the α band.

We estimated, for each of the selected frequency pairs, the degree for each parcel in MEG as the number of connections as a measure of its centrality in the CFC network [80], and averaged degree values over frequency pairs of the same ratio. For SEEG data, we divided again the CFC connectome into the same four laminar depth combinations based on the localization of LF and HF electrode contacts in deep or superficial layers as described in XVII, and then estimated parcel degrees within these sub-connectomes, and averaged, for each ratio and in each group the degree values over frequency pairs. We then estimated the correlation of degree values in MEG CFC connectomes with those in each of the four SEEG groups using a Spearman’s rank correlation test, correcting for multiple comparisons with Benjamini-Hochberg. Fisher’s z-transform was further used to test differences between correlation values within SEEG depth combinations of the same ratio.

#### XIX. Estimation of functional organization of CFC networks

To investigate the functional organization of CFC networks, we identified the brain regions that served predominantly as either LF or HF hubs. We achieved this by using two complimentary approaches: *relative directed degree* and *parcel directionality*.

##### Estimation of relative directed degree

First, in order to be able to compare SEEG and MEG data, all electrode contacts in SEEG, and all parcels of the 200-parcel atlas that had been used so far in MEG, were collapsed to the 148 parcels of the Destrieux atlas [73]. Then, for each LF-HF combination, we estimated for each parcel *p* the graph-theoretic measures *relative in-degree* and *relative out-degree* [80], indicating how often that parcel was either the HF or LF node, respectively, of a CFC connection. The relative in-degree or out-degree of a parcel was defined as the fraction of significant connections *N*_*C*_ for which that parcel was the HF node or LF node, respectively, over the total possible number of possible connections *N*_*C,pot*_. In SEEG data, *N*_*E,pot,S*_ was for each subject *S* the sum of possible, *i.e.*, not excluded, connections from contacts assigned to that parcel to electrodes assigned to other parcels, and *N*_*C,pot*_ was obtained by adding possible CFS or PAC connections over all subjects. In MEG, *N*_*C,pot*_ was simply the number of possible, *i.e.*, not excluded, connections of one parcel to other parcels times the number of subject datasets. Finally, the *relative directed degree* was computed as the difference between relative in-degree and out-degree. Positive values therefore indicated that a parcel was predominantly a HF hub, and a negative value indicated that it was predominantly a LF hub. Relative directed degree values were collapsed over frequency bands for visualization. In order to assess similarity between SEEG and MEG data and between CFS and PAC, we computed the correlation of relative directed degree values across parcels using Spearman’s test (*p* < 0.05).

##### Estimation of parcel directionality

In this approach, we first estimated, for each not-excluded pair of parcels *p*_1_, *p*_2_ of the Destrieux atlas, the LF-HF directionality *Dir*_*LH*_ as the difference in the strengths of CFC connections:

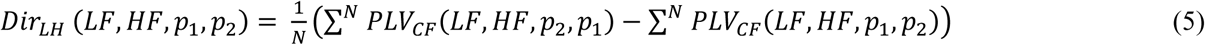

*i.e.*, the total strength of connections where *p*_1_ was the HF node and *p*_2_ the LF node minus the total strength where *p*_2_ was the HF node and *p*_1_ the LF node. *N* was the number of connections between the two parcels, equivalent to *N*_*C,pot*_ defined above. For all pairs (*p*_1_, *p*_2_) in SEEG where *N*_*C*_ < 8, *Dir*_*LH*_ was set to 0.

We then tested, for all frequency pairs, all non-zero values of *Dir*_*LH*_ *(LF,HF,p*_*1*_,*p*_*2*_) for significance value using a permutation test. In each permutation (*N* = 1000), the strengths of all connections *PLV*_*CF*_ *(LF,HF,p*_*1*_,*p*_*2*_*) and PLV*_*CF*_ *(LF,HF,p*_*2*_,*p*_*1*_) between two parcels *p*_*1*_,*p*_*2*_ were pooled and randomly assigned to two groups *G*_*1*_,*G*_*2*_. The permutated directionality *Dir*_*LH*_ value was then computed as

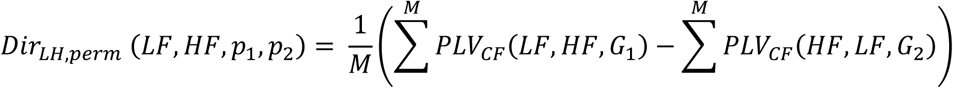

If the genuine *Dir*_*LH*_ value was larger than *Dir*_*LH, perm*_ in 95% of permutations, the directionality of the connection was deemed significant. A significant value of *Dir*_*LH*_ > 0 thus indicated, that between *p*_1_ and *p*_2_, those connections were parcel *p*_*1*_ was the HF node and *p*_*2*_ was the LF node were significantly stronger than those where it was the other around.

The overall directionality 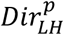 of a parcel *p* was computed as the number of connections with other parcels where its *Dir*_*LH*_ value was positive minus the number of those where it was negative, divided by the total number *N* of possible connections. Thus, similar to the directed relative degree, a positive or negative value of 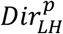 indicated a parcel being a HF hub or LF hub, respectively, in inter-areal CFC. Analogously to relative directed degree, the directionality values were collapsed over frequency bands for visualization, and their similarity between SEEG and MEG and between CFS and PAC was using Spearman’s test (*p* < 0.05).

#### XX. Neuropsychological assessment and correlation of CFC with neuropsychological test results

A subset of the participants of the MEG recordings performed a set of well-validated neuropsychological assessments (*N* = 16–18 for each test). Four tests were used from the Wechsler Adult Intelligence Scale–III (WAIS–III) battery [112]: the Digit Span Forward subtest evaluates verbal short-term memory while the Digit Span Backward and Letter-Number Sequencing (LNS) subtests evaluate verbal working memory, and the Digit Symbol–Coding subtest measures visual psychomotor processing speed. The Trail-Making Test (TMT) [113] is comprised of two parts A and B, of which the TMT-A measures visual scanning and processing speed while the TMT-B measures cognitive flexibility, visual scanning, and processing speed. The Zoo Map Test [114] measures the participant’s planning capability and speed. For the correlation analysis, the test scores were inverted for the TMT and Zoo Map tests, which measure processing speed, so that the higher values were indexing better performance as in the first four tests. To investigate the correlation of the CFC with psychophysical performance, we then computed the correlation of subject’s test scores with their individual graph strength (*GS)* using Spearman’s rank correlation test (*p* < 0.05). To consider the effect of for multiple comparisons across all tests performed, we estimated how many of the total number of the observed significant findings were predicted to be observable by chance. In total, across the eight neuropsychological tests and the CFC frequency pairs, the probability for finding the observed number of significant observations (170) by chance was *p* = 0.0051. Thus we consider the CFS overall to be significantly correlated with neuropsychological tests. The asterisks in Fig 8 indicate the most significant observations (34) exceeding the number of significant observations (136) predicted to be observable by chance at *p* = 0.05. The total number of significant observations for PAC was 45, that is, below the number expected by chance at *p* = 0.5. Thus PAC, at least in such widespread testing, is overall not correlated with neuropsychological test performance.

Data from this study has been deposited in the Dryad depository: https://datadryad.org/stash/dataset/doi:10.5061/dryad.fb240 [115].

## Suppl. Figures

**S1 Fig.**
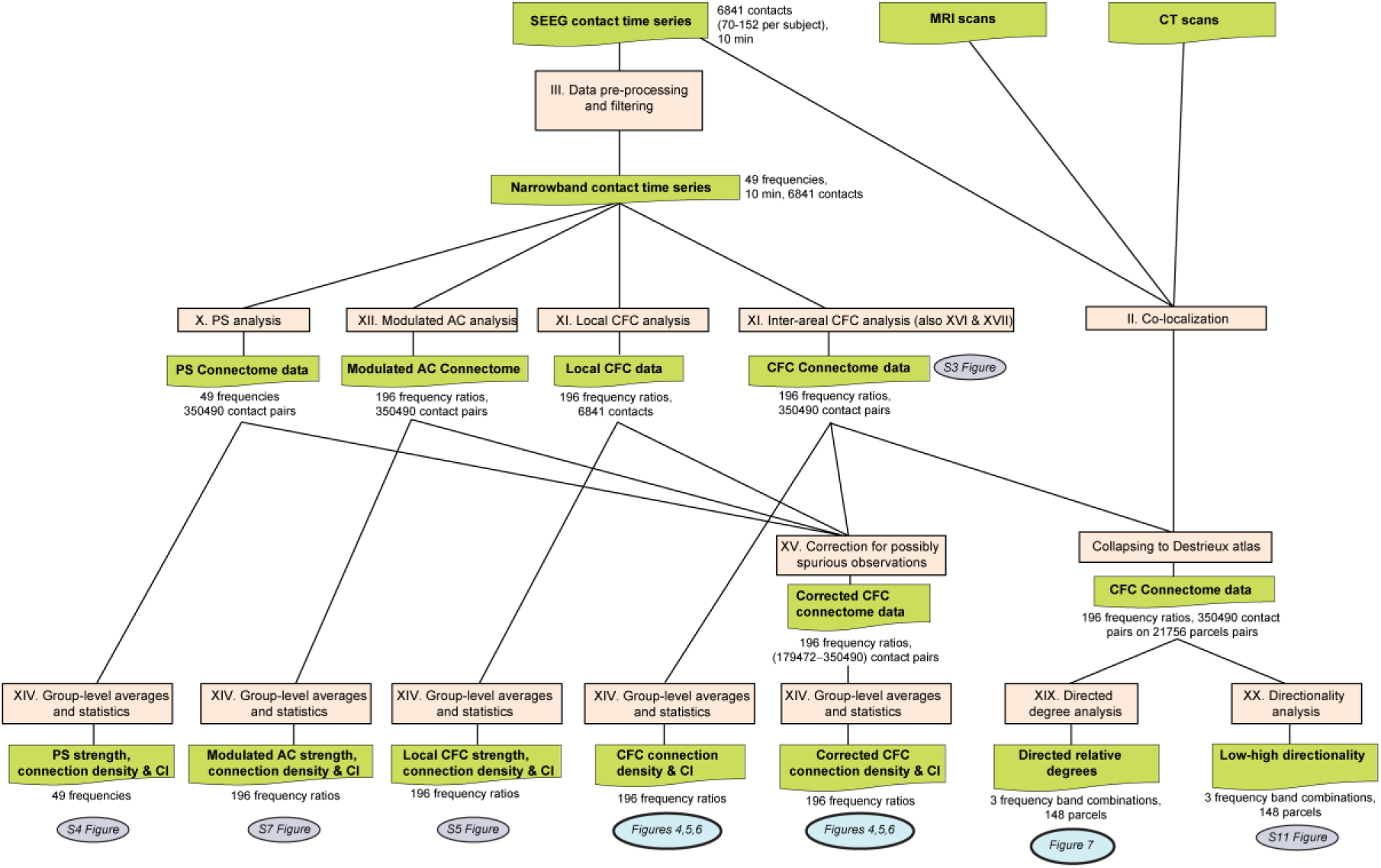
Workflow for SEEG data. Workflow for the processing and analysis of SEEG data, flowing from top to bottom.

**S2 Fig.**
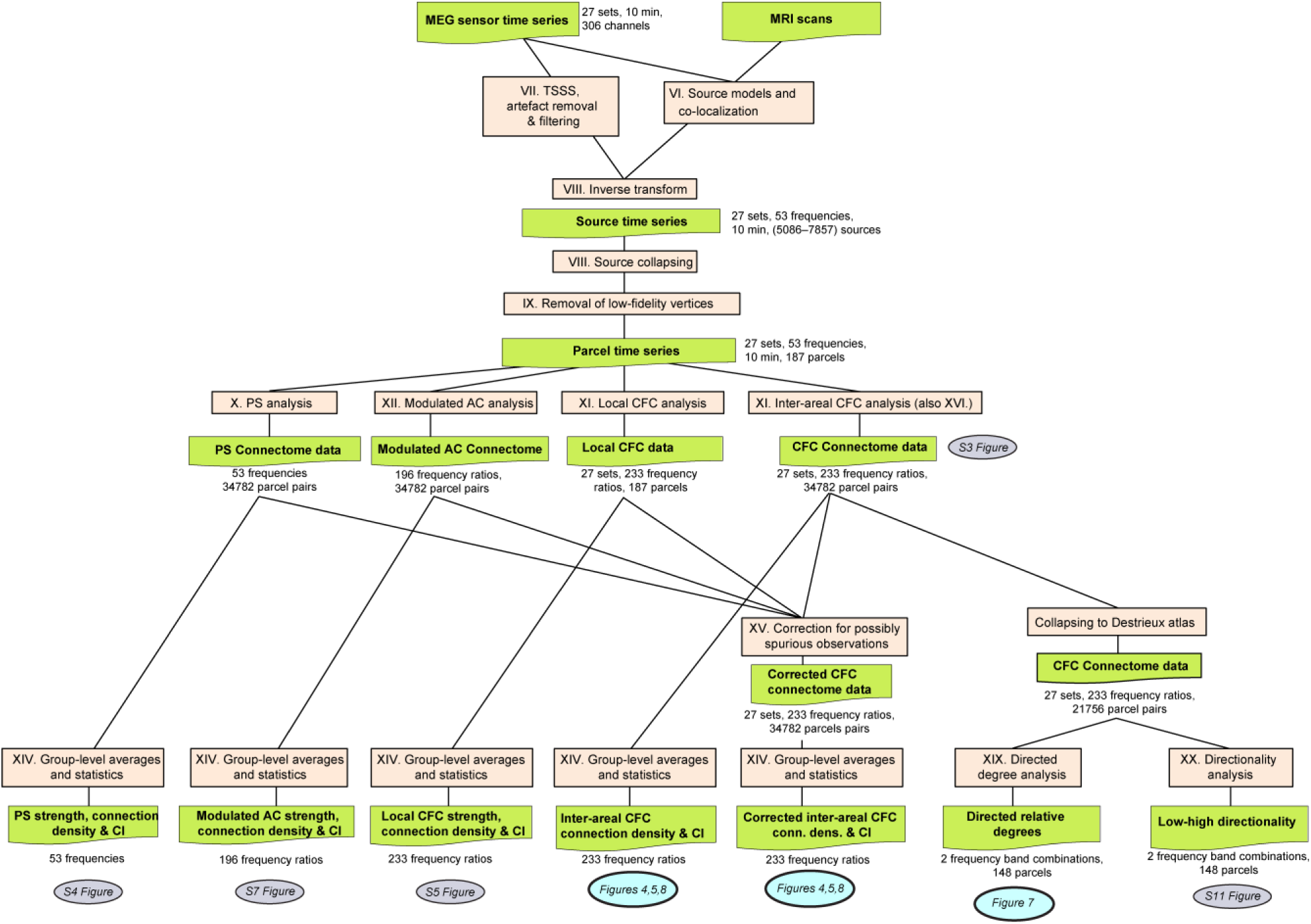
Workflow for MEG data. Workflow for the processing and analysis of MEG data, flowing from top to bottom.

**S3 Fig.**
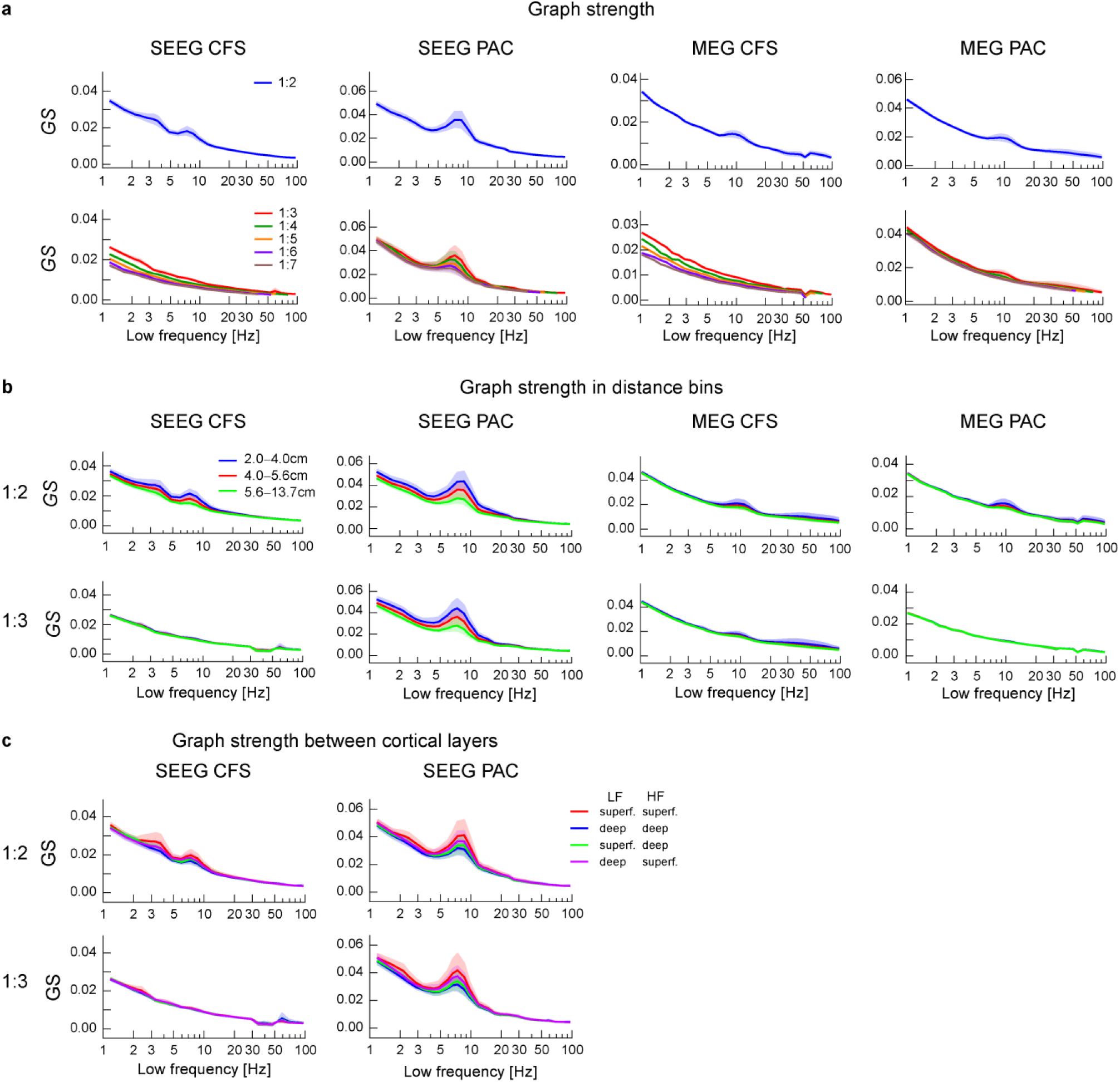
Graph strength of inter-areal CFS and PAC. **a)** The graph strength *GS* (mean of all connections of all subjects) of inter-areal CFC with 95% confidence limits for ratio 1:2 (top row) and ratios 1:3–1:7 (bottom row). **b)** *GS* of inter-areal CFC at ratio 1:2 (top row) and 1:3 (bottom row) in three distance bins, as in Fig 4. **c)** *GS* of inter-areal CFC at ratio 1:2 (top row) and 1:3 (bottom row) in SEEG among different groups of electrode pairs, based on their location in deep or superficial layers, as in Fig 6. Plot data and underlying connectome data are available online [115].

**S4 Fig.**
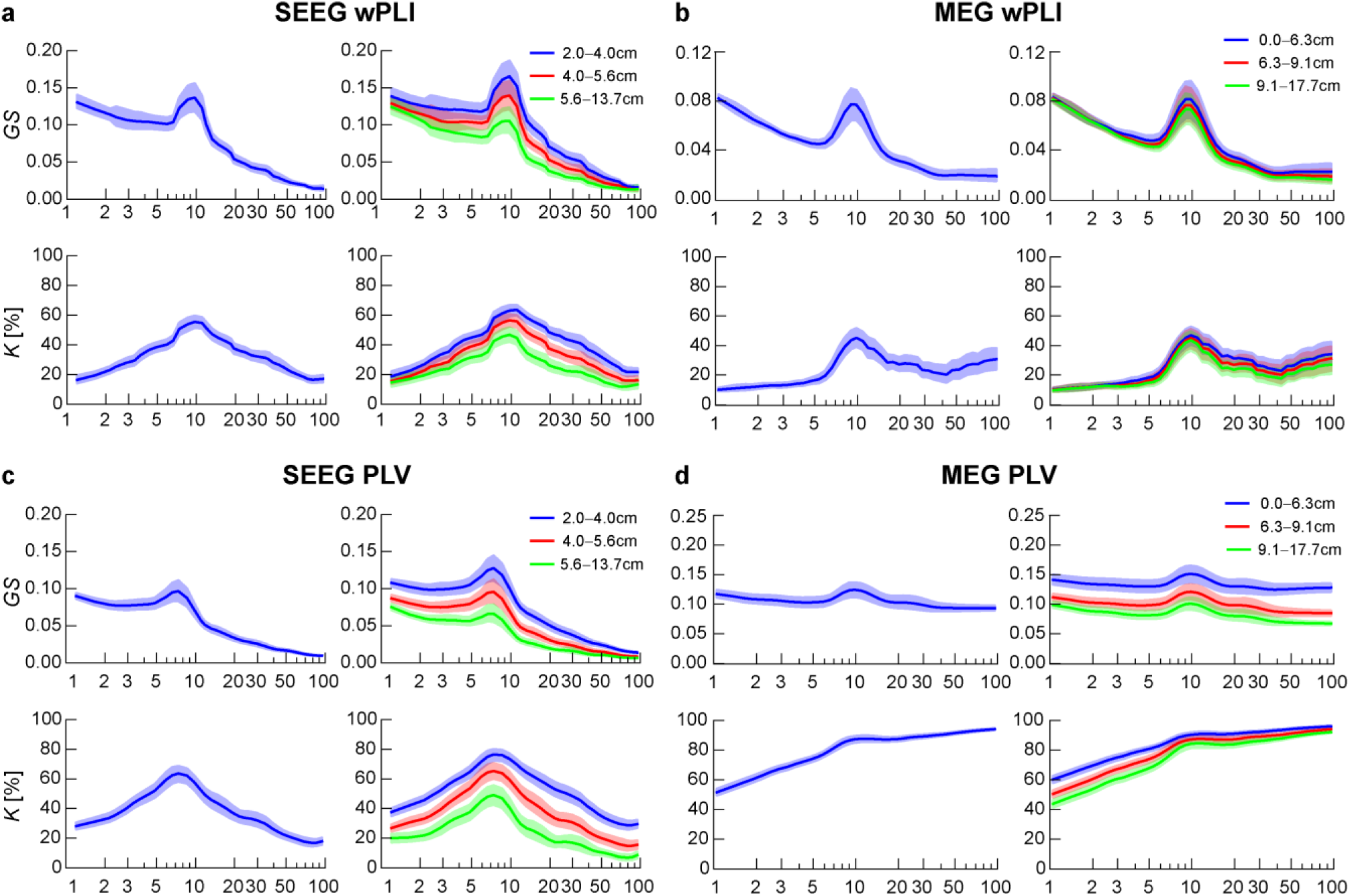
1:1 phase synchrony in SEEG and MEG. **a)** The mean *GS* of 1:1 PS (top) and *K* (bottom) in SEEG data with 95% confidence limits estimated with the weighted Phase-Lag Index (wPLI) for all connections (left) and separately in three distance bins (right). **b)** Same in MEG data. **c-d)** Phase synchrony in SEEG data and MEG data, respectively, estimated with the Phase-Locking Value (PLV). Plot data and underlying connectome data are available online [115].

**S5 Fig.**
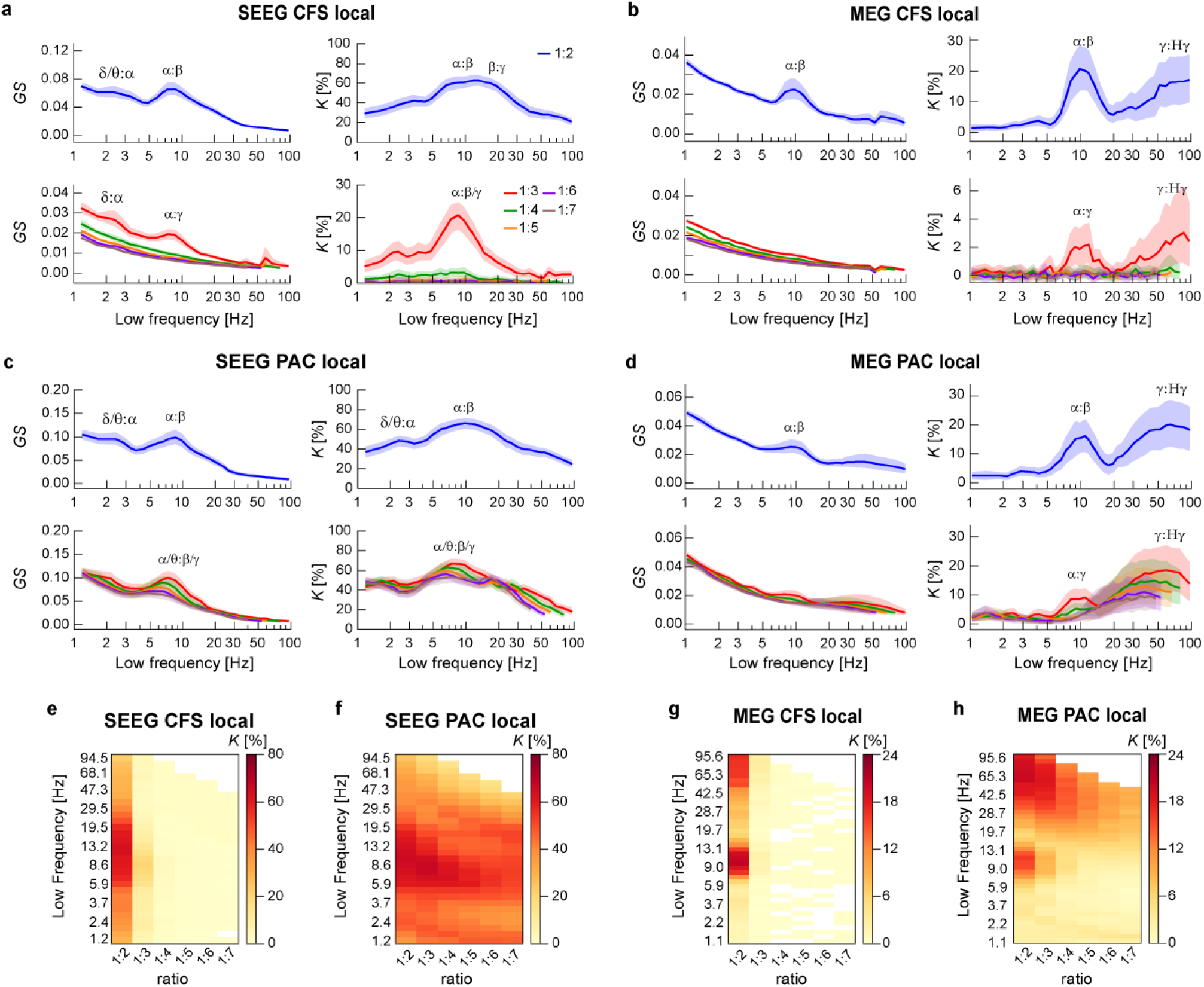
Local CFS and PAC in SEEG and MEG data. **a)** The mean graph strength *GS* and fraction of significant parcels (*K)* with 95% confidence limits (shaded) of local CFS in SEEG data for coupling ratios 1:2 (top) and 1:3 – 1:7 (bottom). Values are shown with 95% confidence limits. **b)** Same for local CFS in MEG data. **c-d)** Same for local PAC in SEEG data and MEG data, respectively. **e)-h)** The same *K* values as shown in a-d plotted in matrices for the whole spectral connectome. Plot data and underlying connectome data are available online [115].

**S6 Fig:**
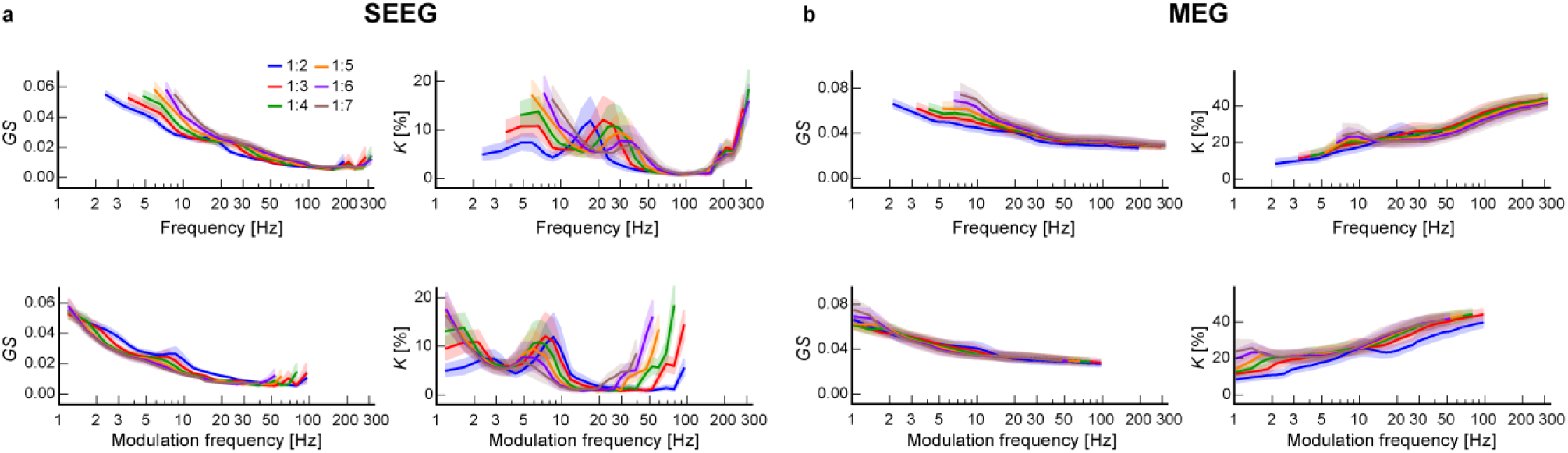
Amplitude envelope correlations. **a)** Graph strength *GS* and *K* of for amplitude envelope correlations in SEEG data for ratios of 1:2 to 1:7. The same data is plotted as a function of the envelope frequency (top row) and the modulating frequency (bottom row). **b)** Same for MEG data. Plot data and underlying connectome data are available online [115].

**S7 Fig.**
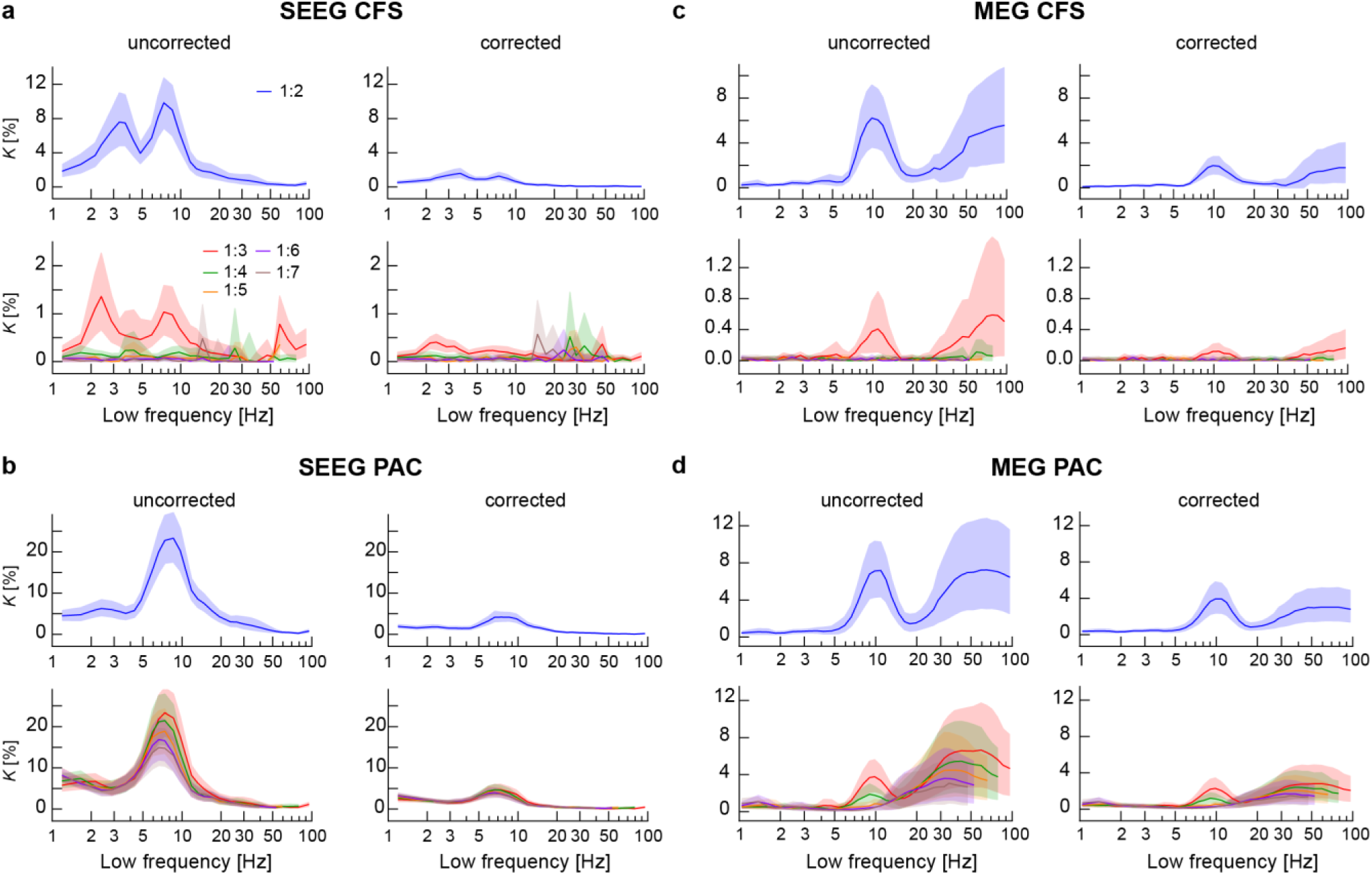
Inter-areal CFS and PAC when PLV is used for removing spurious connections. **a)** Connection density *K* of inter-areal CFS in SEEG data before (left) and after removing possible spurious connections (right) using the PLV as a metric for phase synchrony. **b)** Same for inter-areal PAC in SEEG data. **c)-d)** Same for inter-areal CFS and PAC in MEG data. Plot data and underlying connectome data are available online [115].

**S8 Fig:**
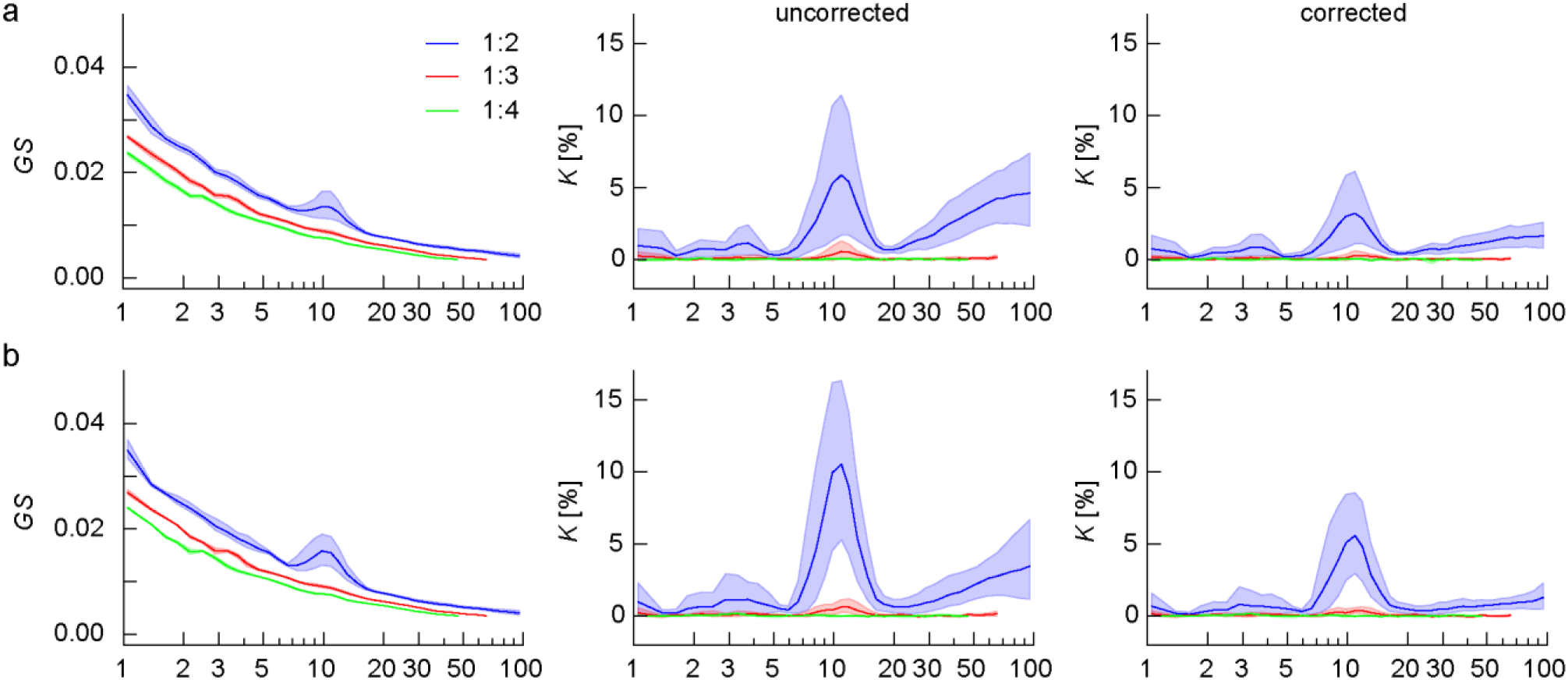
CFS in eyes-open and eyes-closed resting state. **a**) The mean *n*:*m* CFS graph strength (*GS*, left), *K* before correction (middle), and *K* after correction (right, using PLV as PS metric) of inter-areal 1:2 – 1:4 CFS in eyes-open resting state MEG data. **b)** The same in eyes-closed resting state MEG data. Plot data and underlying connectome data are available online [115].

**S9 Fig:**
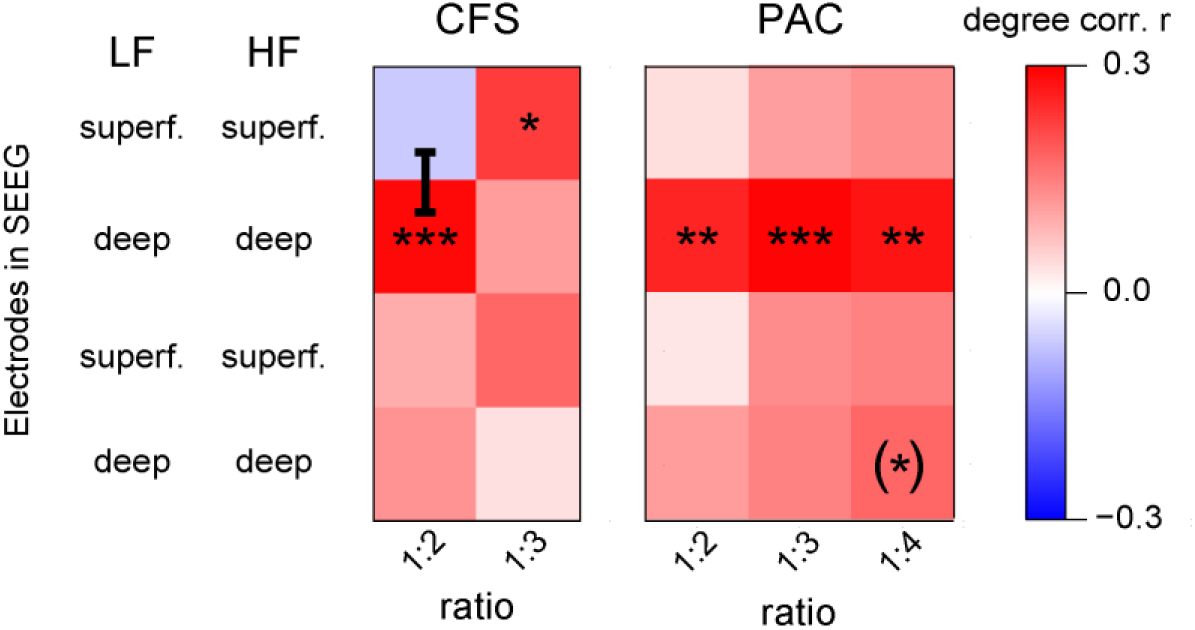
Correlations between MEG data and laminar SEEG data. Correlation of parcel degree values of MEG CFC connectomes with parcel degree values of SEEG CFC connectomes connecting electrodes in either both in more superficial layers, both in deeper layers, from superficial to deep or deep to superficial layers (Spearman’s rank correlation test, ***: p<0.001, **: p<0.01, *: p<0.05, (*): p<0.05, n.s. after correction with Benjamini-Hochberg). The black bar denotes where two correlation values for the same ratio were found to be significantly different by a Fisher’s z-transform test (*z* > 1.96). Plot data and underlying connectome data are available online [115].

**S10 Fig.**
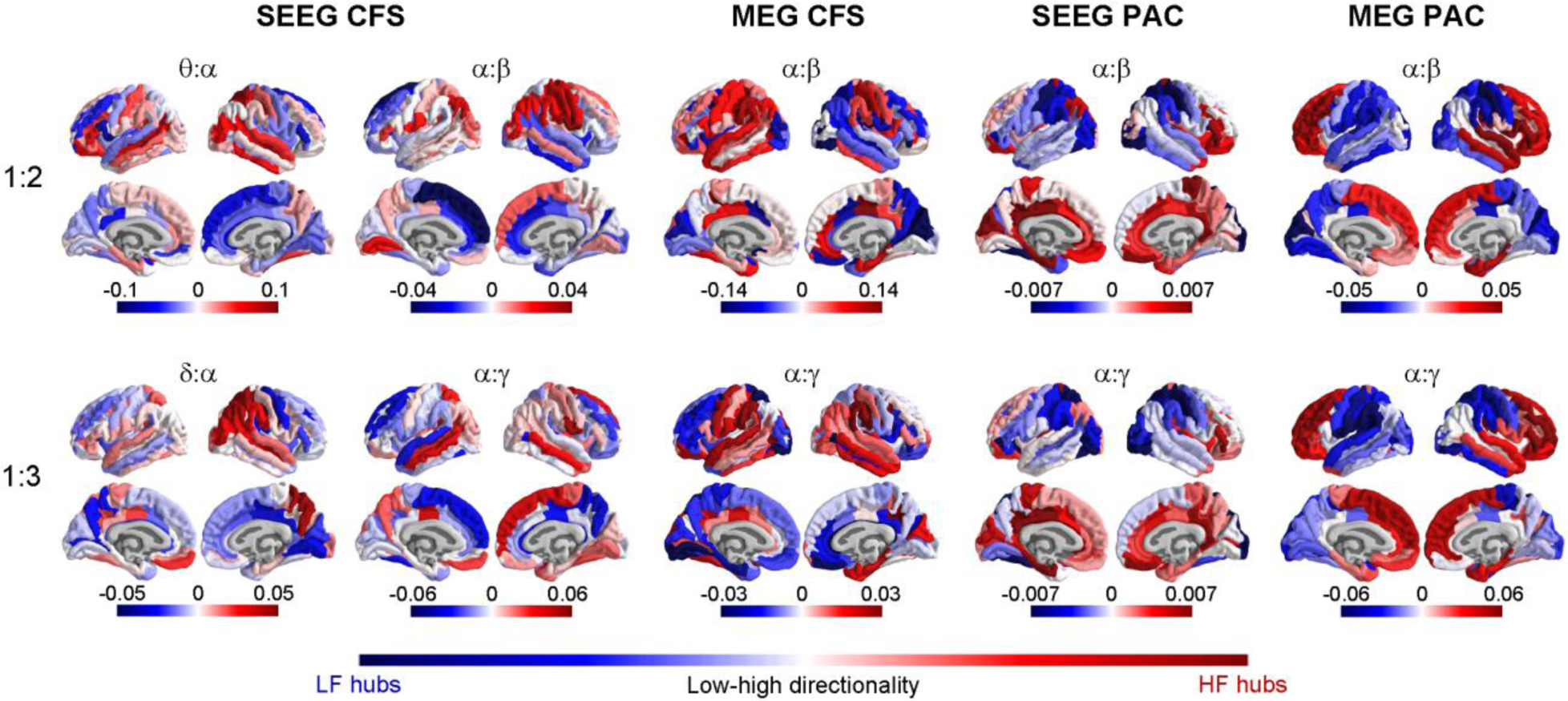
Asymmetric directional connectivity in CFC networks. The organization of CFC networks as measured with directionality between low- and high-frequency hubs. Averaged LF vs. HF directionality values for each parcel. The values indicate whether parcel is a directional hub for LF (blue) or for HF (red) in inter-areal CFC networks. Top row: Directionality for CFS and PAC networks at ratio 1:2 connecting θ:α and α:β frequencies. Bottom row: Directionality for CFS and PAC at ratio 1:3, connecting δ:α and α:γ frequencies. Directional connections of CFS and PAC networks reflect their structure in brain anatomy and show similar opposite directional connections connecting anterior and posterior brain regions, as seen in degree hub analysis (see Fig 7). Plot data and underlying connectome data are available online [115].

**S11 Fig:**
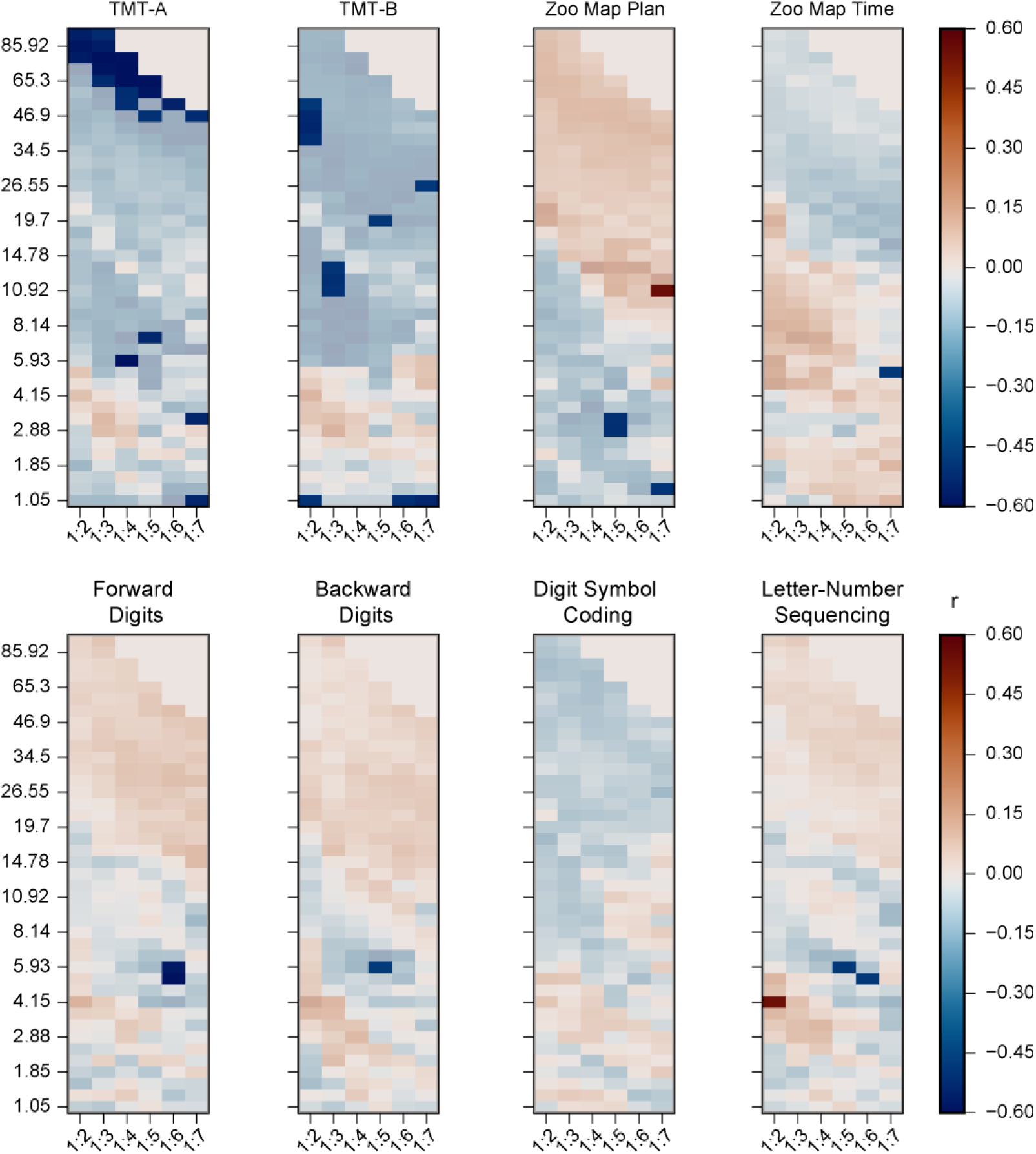
Correlation of PAC with neuropsychological test scores. The correlation or PAC strength in MEG data with neuropsychological test scores (Spearman’s rank correlation test, *p* < 0.05). Red color indicates a positive correlation, while blue indicates a negative correlation, as in Fig 8. Correlations that do not reach statistical significance are masked with lower opacity. No correlations were significant after correcting for multiple comparisons with Benjamini-Hochberg. Plot data and underlying connectome data and neuropsychological data are available online [115].

